# Early-branching cyanobacteria up-regulate superoxide dismutase activity under a simulated early Earth anoxic atmosphere

**DOI:** 10.1101/2024.03.05.583491

**Authors:** Sadia S. Tamanna, Joanne S. Boden, Kimberly M. Kaiser, Nicola Wannicke, Jonas Höring, Patricia Sánchez-Baracaldo, Marcel Deponte, Nicole Frankenberg-Dinkel, Michelle M. Gehringer

## Abstract

The evolution of oxygenic photosynthesis during the Archean (4-2.5 Ga), required the presence of complementary reducing pathways to maintain the cellular redox balance. While the timing of the evolution of superoxide dismutases (SODs), enzymes that convert superoxide to hydrogen peroxide, within the Bacteria and Archaea is not resolved, SODs containing copper and zinc in the reaction centre (CuZnSOD) were the first SODs estimated to appear in photosynthetic cyanobacteria, ≥ 2.93 Ga. Here we analysed the SOD gene expression and activity in the deep branching strain, *Pseudanabaena* sp. PCC7367. It releases more O_2_ and exhibits significantly higher growth rates (p<0.001) and protein and glycogen contents (p<0.05) under anoxic conditions compared to control cultures grown under present oxygen rich atmospheres in low CO_2_ (LC) or high CO_2_ (HC), prompting the question as to whether this correlates to higher cellular SOD activity under anoxic Archean simulated conditions. Expression of *sodB* encoding an iron containing SOD (FeSOD) and *sodC,* encoding a CuZnSOD, strongly correlated with increased extracellular O_2_ levels (p<0.001), while transcription of *sodA*, encoding a manganese containing (MnSOD), correlated to SOD activity during the day (p=0.019), when medium O_2_ concentrations were the highest. Cytosolic SOD activity was significantly higher (p<0.001) in anoxic cultures, two hrs before nightfall compared to oxic growth conditions. Night-time combined *sodABC* transcription in stirred cultures was significantly reduced (p<0.05) under anoxic conditions at elevated CO_2_ levels, as were medium O_2_ levels (p≤0.001), when compared to cultures grown under present-day oxic conditions with low CO_2_. Total cytosolic SOD activity remained comparable, suggesting that the replacement rate of SOD is higher under modern-day conditions than on early Earth. Our data suggest that the early branching cyanobacterium *Pseudanabaena* sp. PCC7367 may have retained ‘ancestral’ features permitting it to thrive in its ecological niche as a benthic mat in shallow water marine environments.

## Introduction

Life on Earth evolved under anoxic, slightly reducing conditions (Fischer & Valentine, 2019; Hamilton, 2019) and was predominated by anaerobic prokaryotes (Hamilton, 2019; Ślesak et al., 2019). Molecular O_2_, a commonly used electron acceptor in aerobic respiration today, was not freely available (Kump, 2008; Lyons et al., 2014). This changed upon the evolution of ancestral photosystems, capable of hydrolysing water and releasing O_2_ during the Mesoarchean, about 3.2-2.8 Ga (Cardona et al., 2019) and their incorporation into early Cyanobacterial species whose crown group emerged between 2.7-2.9 Ga (Boden et al., 2021; Fournier et al., 2021). A few hundred million years passed before the Earth’s atmosphere was enriched with free O_2_ to 1% of present-day levels (Sessions et al., 2009), during a period known as the Great Oxygenation Event (GOE), which began about 2.45 Ga (Bekker et al., 2004; Lyons et al., 2014; Warke et al., 2020). The gradual oxygenation of the environment resulted in the emergence of new enzymes and reaction pathways (Jabłońska & Tawfik, 2021) and is thought to have caused large scale extinction of anaerobic lifeforms unable to cope with uncontrolled oxidation of their cellular components (Case, 2017; Fischer et al., 2016; Fischer & Valentine, 2019; Hamilton, 2019). Free O_2_ would rapidly have been scavenged by reductants such as Fe(II), Mn(II) and ammonia or atmospheric volcanic gases (Ward et al., 2016), however O_2_ levels in microbial mats may have exceeded those of present day atmospheric levels (Herrmann & Gehringer, 2019).

To survive in an increasingly oxygenated environment, organisms had to evolve enzymes to reduce cellular damage from superoxide (O_2_^•-^) (Case, 2017; Hamilton, 2019; Ślesak et al., 2019; Ślesak et al., 2016). Superoxide dismutases (SODs) and superoxide reductases (SORs) both reduce superoxide, but SODs are the only autonomous enzymes known to date that can disproportionate O_2_^•-^ to O_2_ and hydrogen peroxide (H_2_O_2_), which is further converted to water and O_2_ by catalases (CAT) or to water by thiol-, NADH- or cytochrome-dependent peroxidases, thereby enabling cells to maintain their intracellular homeostasis (Case, 2017; Johnson & Hug, 2019). Genetic reconstruction studies have traced the SOD genes to LUCA (the last universal common ancestor) (Ouzounis et al., 2006), suggesting that the ancestral bacterium that gave rise to early Cyanobacteria, may already have been equipped with essential detoxification enzymes (Boden et al., 2021; Johnson & Hug, 2019; Ślesak et al., 2016).

Four different isoforms of SOD exist, based on their metal co-factors; namely CuZnSOD containing copper and zinc, FeSOD containing iron, MnSOD with manganese and NiSOD that has nickel in its active site. Sequence similarity and structure make it difficult to differentiate between MnSOD and FeSOD, suggesting they have a common ancestor (Harada et al., 2021; Johnson & Hug, 2019). However, differentiation of FeSODs and MnSODs is potentially possible based on unique amino acid motifs (Priya et al., 2007). Reduction of Mn^3+^ to Mn^2+^, with its higher midpoint reduction potential, is energetically preferable to the reduction of Fe^3+^ to Fe^2+^, explaining the lower Fenton reactivity of Mn^2+^ and making MnSOD more stable under conditions of oxidative stress (Miller, 2012). Together, the MnSOD/FeSODs are the most widely spread SODs within the Cyanobacteria, with CuZnSODs more rarely encountered (Boden et al., 2021; Harada et al., 2021). NiSODs occur primarily in salt-water strains, mainly in the more recently evolved Picocyanobacteria (Boden et al., 2021; Dupont et al., 2008; Harada et al., 2021). Molecular dating of the four SOD isoforms into the cyanobacterial genomic tree indicates that CuZnSOD was present in the cyanobacterial lineage prior to the GOE, after the Cyanobacteria had diverged from their non-photosynthetic relatives, the Vampirovibrionia (Boden et al., 2021). The genes encoding MnSOD/FeSOD make an appearance after the GOE spreading into new lineages throughout the Proterozoic, while NiSOD appears during the Proterozoic, when Cyanobacteria moved into the open ocean (Boden et al., 2021).

The distribution of SODs within different subcellular compartments of cyanobacteria is dependent on the source of O_2_^•-^ (Fig.1). *Spirulina platensis* expresses a cytosolic FeSOD that is induced under increased salinity and iron availability (Ismaiel et al., 2014). Daytime photosynthesis results in upregulation of *sodB* expression, accompanied by an increase in FeSOD activity in the cytoplasm of *Synechocystis* sp. PCC6803 cultures under normal oxic conditions (Kim & Suh, 2005). Attempts to complement a *Synechocystis* sp. PCC6803 *sodB* deletion mutant with a *sodC* from *Synechococcus* sp. CC9311, revealed that the CuZnSOD was located in the thylakoid membrane and lumen, and was unable to replace FeSOD activity in the cytoplasm (Ke et al., 2014). MnSOD can also be membrane located as observed in *Synechococcus* sp. PCC7942 (Herbert et al., 1992) and *Nostoc* sp. PCC7120 (Li et al., 2002). Therefore, SODs can be directed to membranes, or pass through into compartments (thylakoid or periplasmic space for cyanobacteria), based on the signal peptide encoded on the genome (Russo & Zedler, 2021). Analysis of the leader peptide for MnSOD from *Nostoc* sp. PCC7120, combined with overexpression studies, indicated that several forms of MnSOD were encoded by a single *sodA* gene. These include a full length membrane bound protein, and functional truncated proteins that were located in both the cytoplasm or membrane fractions (Raghavan et al., 2013; Raghavan et al., 2015). FeSOD production increased 6-8-fold during the transition from nitrogen replete to nitrogen depleted conditions in *Nostoc* sp. PCC7120, while MnSOD was principally found on the luminal side of the thylakoid membrane. It was proposed that *Nostoc* sp. PCC7120 could modulate the proteolytic processing of N-terminal signal and linker peptides of membrane-targeted MnSOD in response to nitrogen availability (Raghavan et al., 2015).

**Figure 1:**
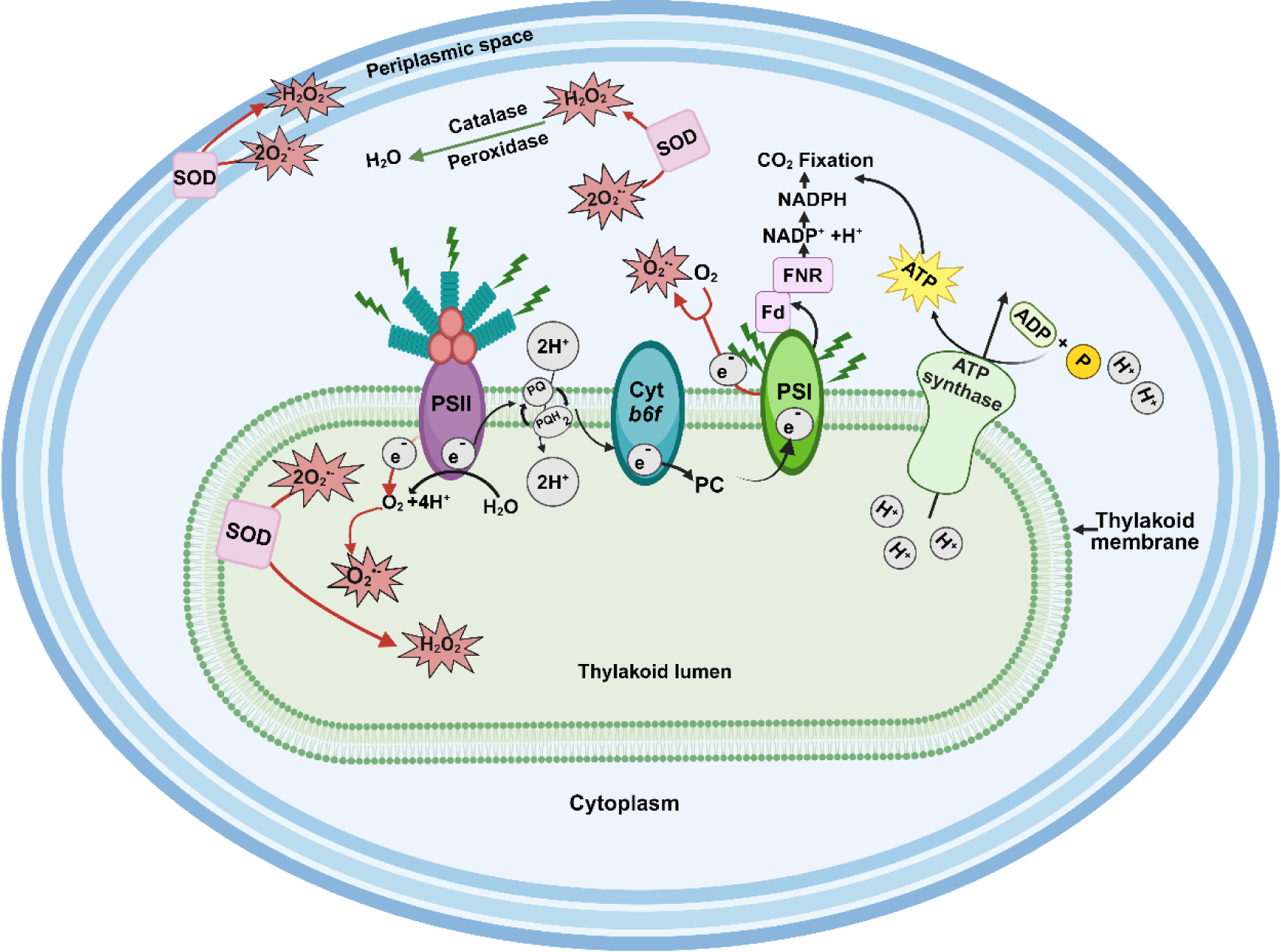
Schematic representation of oxygenic photosynthesis and the generation of superoxide during the day inside a Cyanobacterial cell. The thylakoid membrane contains Photosystem II (PSII) connected to phycobilisomes, Photosystem I (PSI), cytochrome *b6f* (cyt *b6f*), and electron transporters plastoquinone (PQ) and plastocyanin (PC). Phycobilisomes capture light energy and transfer it to PSII resulting in the hydrolysis of water and transfer of electrons to PQ, with the concomitant generation of molecular oxygen (O_2_) and protons (H^+^). PQ then transfers the electrons to PSI via cyt *b6f* and PC, thereby translocating two protons (2H^+^) across the thylakoid membrane, into the lumen. The resulting proton gradient powers the synthesis of adenosine triphosphate (ATP) via ATP synthase. PSI transfers electrons to ferredoxin (Fd), that in turn transfers electrons to ferredoxin NADP+ oxidoreductase (FNR) to generate reduced nicotinamide adenine dinucleotide phosphate (NADPH). Both NADPH and ATP are used to power cellular processes, such as CO_2_ fixation (Mullineaux, 2014). Increased light exposure may result in excess electrons being fed into the electron transport chain, resulting in the generation of superoxide (O_2_^•–^) (Latifi *et al.,* 2009). Dismutation of O_2_^•–^ to hydrogen peroxide (H_2_O_2_) potentially occurs in different species via superoxide dimutases (SODs), in the cytoplasm, thylakoid lumen or periplasmic space (Herbert et al., 1992; Li et al., 2002; Napoli et al., 2021; Raghavan et al., 2013; Raghavan et al., 2015). Image generated in Biorender ©.

Export of MnSOD to the thylakoid lumen or periplasmic space is confirmed supported by sequence analysis of one of the two MnSODs encoded by *Chroococcidiopsis* sp. CCMEE 029. A signal peptide for the TAT signal transduction system was identified for one of the MnSODs (SodA2.1), suggesting that it is localised in the periplasmic space and/ or the thylakoid lumen (Napoli et al., 2021). The second MnSOD (SodA2.2) and the CuZnSOD (SodC) carried no signal-peptide and are presumably present in the cytoplasm. Expression levels of all three SOD genes were significantly elevated after 60 min desiccation in the dark (Napoli et al., 2021). In summary, the type, number and exact location of SOD isoforms may depend on the environment, species evolution and functionality. Yet, no research on SOD gene expression and activity in early-branching cyanobacteria, such as *Pseudanabaena* spp., has been conducted.

Molecular clock analyses have identified a deep branching clade of cyanobacterial species that can be traced back to the late Archean, prior to the great Oxygenation Event (Boden et al., 2021; Jahodářová et al., 2018; Sánchez-Baracaldo, 2015; Sánchez-Baracaldo et al., 2017). It includes several *Pseudanabaena* spp., and net O_2_ production rates by one of these early branching marine strains, *Pseudanabaena* sp. PCC7367, are significantly higher in cultures grown anoxically at 0.2% atmospheric CO_2_ (Herrmann et al., 2021). Control cultures grown under present-day levels of CO_2_ and O_2_, or controls supplemented to 0.2% CO_2_ demonstrated significantly lower rates of O_2_ release per chlorophyll a content. This raises the question as to the potential of host SODs inactivating the potentially increased levels of O_2_^•-^ in cyanobacteria growing in a simulated oxygen free, early Earth atmosphere. *Pseudanabena* sp. PCC7367 encodes three putative SODs, namely MnSOD (*sodA*), FeSOD (*sodB*), and *sodC,* encoding CuZnSOD (Boden et al., 2021; Harada et al., 2021). This study investigates the expression and activity of these SODs, over 24 hrs under a simulated high CO_2,_ anoxic atmosphere to assess the role of atmospheric O_2_ on cyanobacterial growth. Altogether, our data provides insight into diurnal expression and activity of SODs in a deep branching Cyanobacterium under an anoxic, early Earth simulation compared to present-day oxygen rich conditions.

## Materials and methods

### Culture conditions

*Pseudanabaena* sp. PCC7367 was purchased from the Pasteur Culture Collection (Paris, France) and maintained under normal present atmospheric levels (PAL) of 0.04% CO_2_ (low CO_2_ - LC), under photosynthetically active photon flux (PPFD) of 20 μmol photons m^-2^. s^-1^, 65% humidity and a (16:8) day:night cycle in artificial salt medium, ASNIII (Hermann *et al.,* 2021) (25). The experimental cultures used in this study were similarly maintained for 6 months under their respective atmospheric conditions of PAL (LC) or PAL supplemented to 0,2% CO_2_ (HC) in a Percival E22 growth chamber (CLF Plant Climatics, Germany) illuminated with Radium NL 18W/840 Spectralux Plus white light bulbs (Germany). Similarly, cultures were grown under anoxic Archean simulated conditions in N_2_ gas containing 0.2% CO_2_ (Archean) in an anaerobic workstation (MEGA-4, GS Glovebox, Germany) fitted with Cree CXB1816 LED lights (USA). We ensured the light spectra and light intensities were the same under all three growth conditions, using an optical multichannel analyzer (USB2000^+^ Ocean Optics, Germany).

Comparative growth curves of acclimated cultures were initiated at an initial Chl a concentration of 0.04 μg. mL^-1^ of chlorophyll a (Chl a) in 500 ml ASNIII medium, in triplicate, in large Fernbach flasks for each of the three atmospheric conditions. The Chl a content was measured every second workday, with samples for the assessment of cellular carotenoid, protein, and glycogen content collected in parallel, for a total of 28 days. For SOD transcriptional and activity analyses, cells were harvested at a similar Chl a concentration (∼2 μg. mL^-1^) during the late exponential phase to maximize RNA and protein yields, on days 8-9 under Archean atmospheric conditions, days 12-13 for the HC and days 13-14 for the LC culture conditions.

The total number of cells per millilitre was determined using a Neubauer counting chamber (Carvalho *et al.,* 2021) and used to calculate cellular SOD activity. The cell count was performed for each biological triplicate on day 9 for the cultures grown under an Archean simulated atmosphere, day 12 for HC and day 13 for LC grown cultures, the same day on which the protein and RNA samples were taken.

### Chlorophyll a (Chl *a*) and carotenoid extraction

A 2 mL volume of culture was collected in 2 mL brown tubes, centrifuged (5 min, 10000 RCF, Hermle LaborTechnik GmbH - Z 233 M-2 Microliter Centrifuge) and the cell pellet drained. Around 100 μg (a small spatula full) of 0.1 mm silica beads (BioSpec, USA) and 1.5 mL of neutralized 90 % MeOH were added to the drained pellet, followed by disruption (FastPrep bead beater FP 120, Thermo Electron Corporation) at 6.5 m. sec^-1^ speed for 45 seconds (twice) and incubation overnight in the dark at 4 °C. The next day, the lysate was vortexed, centrifuged (15 min at 10000 RCF), and the absorbance of the supernatant measured at 665 nm (Chl a) and 470 nm (carotenoid), using a spectrophotometer (Hellma, Agilent 8453, China). Chl a (Meeks & Castenholz, 1971) and carotenoid (Wellburn, 1994) content was calculated (Herrmann et al., 2021). The growth rate was determined from the Chl a growth curve for days 0 to day 12.

### Protein and glycogen quantification

The protein and glycogen content of cells provide insight into the growth of the culture and its nitrogen and carbohydrate resources (Herrmann & Gehringer, 2019; Klotz et al., 2016). A cell lysate was prepared from 2 mL pelleted culture material (12000 RCF for 5 min) that was resuspended in 1 mL of lysis buffer (5 mM NH_4_SO_4_, 2 mM DTT, 1 mM MgCl_2_, 20 mM KH_2_PO_4_ (pH 8.5)) with ∼100 μg (a small spatula full) of 0.1 mm silica beads (BioSpec, USA), followed by mechanic disruption as above. The samples were additionally freeze/thawed in liquid nitrogen before and after each round of bead beating. To pellet cell debris, the cell extracts were centrifuged at 13000 RCF for 15 min (HERMLE, Z233 M-2, Germany) and the supernatant transferred to a fresh 1.5 mL tube and stored at −20 °C. These cell lysates were used for both protein and glycogen determinations.

The protein content of the biomass was assessed using the Bradford assay. A 1:2 serial dilution series of an Albumin Fraction V (Sigma-Aldrich, USA) stock solution of 2 mg. mL^-1^ was made ranging from 0.488 μg. mL^-1^ to 500 μg. mL^-1^, in lysis buffer. For protein determination, a 50 μL volume of blank, standard dilution or sample was added into the wells of a 96-well clear bottom plate and 250 μL of Bradford reagent (Merck, Darmstadt, Germany) was added. The plate was incubated at room temperature for 10 min after which the absorbance at 595 nm was determined (Multiskan FC, Thermo Fisher Scientific). Experimental protein concentrations were read off the standard curve (Herrmann & Gehringer, 2019). R^2^ values ranged between 0.95 to 0.99.

In order to measure the glycogen content of the cells, a 1:2 serial dilution series of a stock solution of oyster glycogen (Sigma, Germany) of 2 mg. mL^-1^ was made ranging from 0.488 μg.mL^-1^ to 500 μg.mL^-1^ in the lysis buffer. A volume of 200 μL of standard dilution, sample or blank was transferred into a 2 mL reaction tube and a volume of 500 μL ice-cold anthrone (Sigma-Aldrich, Germany) reagent (2% w/v in 98% sulphuric acid) was added and then incubated for 30 min at 80°C. Afterwards, 250 μL of each sample was transferred into individual wells of a clear bottomed 96-well plate and the absorbance read at 620 nm (Multiskan FC, Thermo Fisher Scientific). Experimental glycogen concentrations were calculated off the standard curve (Herrmann & Gehringer, 2019). R^2^ values ranged between 0.98 to 0.99.

### Quantification of oxygen medium levels

Oxygen accumulation both within and outside the cell, influences SOD expression levels (Kim & Suh, 2005). Oxygen levels in stationary cultures were measured over 24 hrs to determine the time points for assessing expression and activity of SOD in *Pseudanabaena* sp. PCC7367, grown under the three different atmospheres investigated. The O_2_ levels were initially recorded in cultures without agitation to mimic the proposed shallow water marine oxygen oases identified in the Archean fossil record (Catling & Zahnle, 2020; Riding et al., 2014). Additionally, the cultures were stirred to ensure maximal O_2_ release from the medium during the dark phase, to identify a time point with the lowest possible SOD activity. Levels of dissolved O_2_ (µM. L^-1^) in the culture media of *Pseudanabaena* sp. PCC 7367 (n=3) was measured (Robust Oxygen Probes OXROB10, attached to the Firesting-O2, Pyroscience, Germany) over a full diurnal cycle under each atmospheric condition investigated. Cultures were then gently stirred and the dissolved O_2_ levels again recorded.

### SOD protein characterisation and structural prediction *Pseudanabaena* sp. PCC7367

The three SODs encoded within *Pseudanabena* sp. PCC7367 (Boden et al., 2021; Harada et al., 2021) were investigated with respect to their potential membrane binding and signalling domains. To identify whether transmembrane domains existed and where, the amino acid sequences of each SOD were subjected to DeepTMHMM (Hallgren et al., 2022).

The 3D models of the SODs encoded by *Pseudanabaena* sp. PCC7367 (Boden et al., 2021; Harada et al., 2021) were generated on the Phyre2 model server (Lawrence A. Kelley et al., 2015) and visualized using the Swiss-PDB viewer (http://www.expasy.org/spdbv/) (Guex et al., 2009). Alphafold 2 (Jumper et al., 2021; Varadi et al., 2021) was used to generate the protein structures of the CuZnSOD, MnSOD and FeDOS of *Pseudanabaena* sp. 7367. Final images were coloured for confidence in the secondary structure predictions using PyMOL 2.3 (DeLano, 2020).

### Genetic potential of SOD associated genes in *Pseudanabaena* sp. PCC7367

The KEGG Database (Kanehisa et al., 2022) was searched for annotated superoxide reductases (SORs), peroxidases/peroxiredoxins and catalase genes encoded on the *Pseudanabaena* sp. PCC7367 genome (Shih et al., 2013). The genome of *Pseudanabaena* sp. PCC 7367 (NC_019701.1) was additionally screened, using tBLASTn (Altschul, 1991), for the presence of SORs using the sequences from the Archaea, *Pyrococcus furiosus* DSM 3638 (PF1281) and *Desulfovibrio vulgaris* DP4 (Dvul_0204), obtained from the Superoxide Reductase Gene Ontology Database (SORGOdp) (Lucchetti-Miganeh et al., 2011), as well as the catalase gene from *E. coli* K12 MG1655 (YP_025308). Furthermore, to ensure the complementary metal cofactors were accessible for SOD isoform activities, the genome of *Pseudanabaena* sp. PCC7367 was screened for the presence of the Mn(II) transporters, MntABC (Bartsevich & Pakrasi, 1995), Zn(II) transporters, ZnuABC and Cu(II) transporters (CtaA and PacS) (Sharon et al., 2014), using the characterized protein sequences from *Synechocystis* sp. PCC6803 (NC_000911.1).

### Detection of SOD gene expression levels

A set of primers were designed to target the *sodA* (MnSOD), *sodB* (FeSOD) and *sodC* (CuZnSOD) genes identified in *Pseudanabaena* sp. PCC7367 (Supp. Table 1), as well as the housekeeping RNA polymerase beta subunit gene, *rpoC1* (Alexova et al., 2011; Enzingmüller-Bleyl et al., 2022) using the NCBI Primer-BLAST Software (Ye et al., 2012). To avoid self-complementarity, scores generated by PCR primer inspector (www.molbiotools.com/primerinspector.php) and NetPrimer (www.premierbiosoft.com/netprimer/) were considered. Primers were validated by sequencing of the PCR products as well as determining their binding efficiencies to both genomic DNA (gDNA) and copy DNA (cDNA) generated from total RNA extractions (Supplementary Table 2).

Samples for RNA extraction were taken in late exponential phase on days 8-9 for cultures grown under anoxic conditions, days 12-13 for cultures cultured under an HC conditions and days 13-14 for LC atmospheric conditions, with Chl a concentrations of ∼2 μg. mL^-1^, that had similar protein and glycogen concentrations (Supplementary Fig. 1). Based on the observed dissolved O_2_ levels in stationary cultures (Supplementary Fig. 2), samples were collected two hrs after the lights went on, two hrs before they went off (14 hrs: at maximum levels of dissolved O_2_ in the medium), two hrs after the lights went out (18 hrs) and two hrs prior to them being switched on again (22 hrs). The following evening, an additional sample (19 hrs) was taken, three hrs after darkness, from the stirred cultures, for the assessment of the baseline SOD expression levels at minimum oxygen levels in the dark.

A 90 mL culture volume was added to 10 mL of ice-cold stop solution (5 % Roti®-Aqua-Phenol (ROTH, Germany) in p.A. ethanol) at each time point. The cells were harvested by centrifugation (12000 RCF for 10 min, Hermle Z 513K, Germany) and the pellet weighed. RNA was extracted using the NucleoSpin® RNA Plant Kit (MACHEREY-NAGEL, Germany) according to the manufacturer’s instructions. The initial lysis step was modified to increase the yield of RNA as follows. An approximately 200 mg pellet was resuspended in 350 μL of RA1 solution, then transferred to sterile, RNAse-free 2 mL tube containing around 100 μg of 0.1 mm silica beads (Biospec) and 3.5 μL of β-mercaptoethanol (ROTH, Germany). Cells were subjected to two cycles of a rapid freeze / thaw lysis step in liquid nitrogen followed by mechanical disruption (FastPrep) for 45 seconds at 6.5 speed. The cell debris was removed by centrifugation (1 minute, 11,000 RCF and the supernatant used for further RNA purification following the manufacturer’s instructions. RNA was quantified using a NanoDrop® Lite Spectrophotometer (ThermoFisher Scientific), while the quality was checked by agarose gel electrophoreses on a 1% w/v Tris-acetate-EDTA (TAE) gel (Supplementary Fig. 4). The purity was determined by PCR using the housekeeping gene primer pair to ensure no DNA remained. RNA was subjected to a repeat DNA digestion if a positive PCR was observed. A 10 μL volume of 10-fold DNAse I reaction buffer (NEW ENGLAND BioLabs, catalogue-B0303S), 5 units (2.5 μL) of RNase free DNAse I (NEW ENGLAND BioLabs, 2000 u. mL^-1^) and 100 μL of nuclease free water was added for every 10 μg of RNA, and incubated for 30 min at 37 °C. After DNAse treatment, the quantity and quality of RNA was again checked as previously described. Complementary DNA (cDNA) was reverse transcribed from ∼1 μg high quality RNA using the ProtoScript^®^ II First Strand cDNA Synthesis Kit (NEW ENGLAND BioLabs, Germany). Newly synthesized cDNA was purified using a the NucleoSpin® Gel and PCR Clean-up kit (Germany) and quantified using a NanoDrop® Lite Spectrophotometer. A test PCR using the control primer pair was conducted to confirm successful cDNA synthesis.

Expression of each gene ie: *sodA, sodB, sodC* and *rpoC1* was assessed using the 2x iTaq Universal SYBR Green Supermix (Bio-Rad, USA) and the respective primer pairs. An eight μL volume of master mix containing one μM of forward and reverse primer was aliquoted into the 96-well PCR plate (STARLAB, Germany) for each reaction, and ten ng cDNA in two μL was added. For the no template controls (NTCs), two μL of ddH_2_O was added instead of template. The reaction volumes were mixed by pipetting, the plate was sealed with plastic adhesive foil (Bio-Budget, Germany) and labelled. Three technical replicates were conducted for each reaction, on two different days with an initial incubation of 10 min. at 50 °C to activate the polymerase, followed by a single denaturing step of 5 min. at 95 °C and then 40 cycles of 10 sec. at 95 °C, 20 sec. at 55 °C and 10 sec. at 72 °C, and a final elongation step of five min. at 72 °C. The relative expression of the SOD genes relative to *rpoC1* housekeeping gene was calculated as described in the Supplementary Text.

### Assessment of SOD activity

Protein was extracted from *Pseudanabaena* sp. PCC 7367 cultures during the late exponential phase, at the same time points and similar Chl *a* concentration as the transcriptional analysis (Supplementary Fig. 1). The cell pellet from a 50 mL culture volume was obtained by centrifugation at 8000 RCF for 10 min. (Hermle Z 513K, Germany). The pellet was resuspended in two mL of freshly prepared lysis buffer (5 mM NH_4_SO_4_, 2 mM DTT, 1 mM MgCl_2_, 20 mM KH_2_PO_4_ (pH 8.5)) and sonicated on ice for six cycles of one min., with a break between every minute, at 4 ^°^C, 130W, 20 kHz and 50 % amplitude using an ultrasonicate homogenizer (UV 220, Bandelin, Germany). The sonicated extract was centrifuged (Sorvall Lynx 6000, Thermo-Fischer, Germany) at 43 185 RCF, at 4 ^°^C (60 min.). The supernatant was carefully transferred into a new two mL tube and kept at 4 ^°^C before performing the colorimetric SOD enzyme activity assay. The concentration of the extracted protein was measured using the Bradford assay as previously described.

The total SOD activity of soluble cytosolic proteins was determined using a microtiter plate enzyme inhibition assay (Peskin & Winterbourn, 2017). The assay is based on the reduction of the water-soluble tetrazolium salt (WST-1) with a superoxide anion, producing a water-soluble formazan dye that can be calorimetrically quantified by measuring the absorbance between 410 nm to 450 nm. The superoxide anion is produced through the oxidation of hypoxanthine by xanthine oxidase. SOD inhibits the formation of formazan by catalysing the dismutation of the superoxide anions into hydrogen peroxide and molecular oxygen, thereby decreasing the reduction of WST-1 and hence the absorbance. An assay buffer was prepared with 100 mM of sodium phosphate (pH 8.0), 0.1 mM of diethylenetriamine-Penta acetic acid (Sigma-Aldrich, Germany) and 0.1 mM of hypoxanthine (Sigma-Aldrich, Germany). A stock solution of ten mM hypoxanthine (Sigma-Aldrich) solution was prepared in dimethyl sulfoxide (DMSO, ROTH, Germany) and then diluted as required. Ten ml of a ten mM solution of WST-1 (Dojindo Molecular Technologies, USA) was prepared and wrapped in aluminium foil to protect it from light. A two mg. mL^-1^ solution of catalase (Sigma-Aldrich) was dissolved in phosphate buffer pH7, and stored at 4 ^0^C. Bovine CuZnSOD (Sigma-Aldrich) was used to generate a standard curve or relative SOD activity as follows: a (1:2) dilution series of Bovine CuZnSOD was prepared ranging from 0.0488 μg. mL^-1^ to 100 μg. mL^-1^ in lysis buffer. A one µl volume of blank, standard dilution or sample was added to wells of a 96-well plate, in duplicate, at 24 °C. Afterwards, 0.2 mL of assay buffer containing sufficient xanthinin oxidase (Peskin & Winterbourn, 2017) is added into each well. The 96 well plate was immediately placed into the plate reader (Infinite f200 PRO, TECAN) and shaken for five seconds, after which the absorbance was measured at 415 nm for 5 minutes. The amount of CuZnSOD added in the assay (X) was plotted against the calculated percentage of the inhibition of WST-1 reduction (Y), from which the amount of activity relative to Bovine CuZnSOD were determined. The R^2^ values fell between 0.96 to 0.98 (Supplementary Fig. 5).

### Statistics

Statistics were carried out using tools available in SigmaPlot 13.0 (Systat Software Inc.). The data sets were tested for homogeneity of variance, using Levene’s test, and for normality, using the Kolmogorov-Smirnov test. Differences in the growth associated variables of chlorophyll a, carotenoid, protein and glycogen content, as well as media O_2_ concentration were assessed using repeated measure One Way Analysis of Variance/on ranks, followed by Tukey’s honest significant difference (HSD)/ Holm-Sidak method as post-hoc test.

Two Way Repeated Measures ANOVA (Two Factor Repetition) with treatment and time as factors for parameters sampled during the 24 h sampling cycle, was applied. Those included SOD gene expression levels (*sodA* (MnSOD), *sodB* (FeSOD), *sodC* (CuZnSOD), *sodABC* (total SOD expression) and SOD enzyme activity, as well as concentration of dissolved O_2_. In case of significant differences in parameters caused by environmental conditions, Duncan’s Method was used as a post-hoc test to identify diverging groups. Moreover, Pearson’s correlation was then conducted to identify the relationship between variables of 24 h sampling.

## Results

Comparative growth curves were set up under the three defined atmospheric conditions, to assess which best supported *Pseudanabaena* sp. PCC7367 growth. This permitted the identification of comparative sampling time points for quantification of SOD transcription, and activity, over a full diurnal cycle.

### Growth characterization

Growth rates were determined from the Chl a growth curves generated for *Pseudanabaena* sp. PCC 7367 grown under the Archean simulated anoxic atmosphere, and the HC and LC oxygen-rich atmospheric conditions (Fig. 2A; Supplementary Table 6). *Pseudanabaena* sp. PCC 7367, grown under an anoxic atmosphere exhibited a significantly higher growth rate compared to those cultures grown under present day atmospheric oxygen levels at LC (p<0.001) and HC (p<0.001) (Fig. 2B; Supplementary Table 6). Carotenoid levels were also significantly higher in anoxic grown cultures, compared to cultures grown under LC (p=0.001) conditions (Supplementary Fig. 3A; Supplementary Table 6). However, the Chl a:carotenoid ratio was similar on the days of sampling (Arrows Fig. 2B). Glycogen levels were significantly raised under anoxic growth conditions (Supplementary Fig. 1B; Supplementary Table 6) compared to LC (p=0.004) and HC (p=0.035) atmospheres (Tukey Test: all pairwise multiple comparison procedures). Protein levels were significantly higher in culture material from Archean grown cultures, than biomass from LC (p=0.002) or HC (p=0.029) growth conditions (Supplementary Fig. 1C; Supplementary Table 6). In summary, *Pseudanabaena* PCC7367 grown under anoxic conditions had a significantly higher growth rate than cultures grown under oxic conditions. The significantly higher levels of glycogen and protein accumulation in stationary phase cultures point to increased cell vitality, permitting the accumulation of excess carbon, while maintaining the carbon to nitrogen balance.

**Figure 2.**
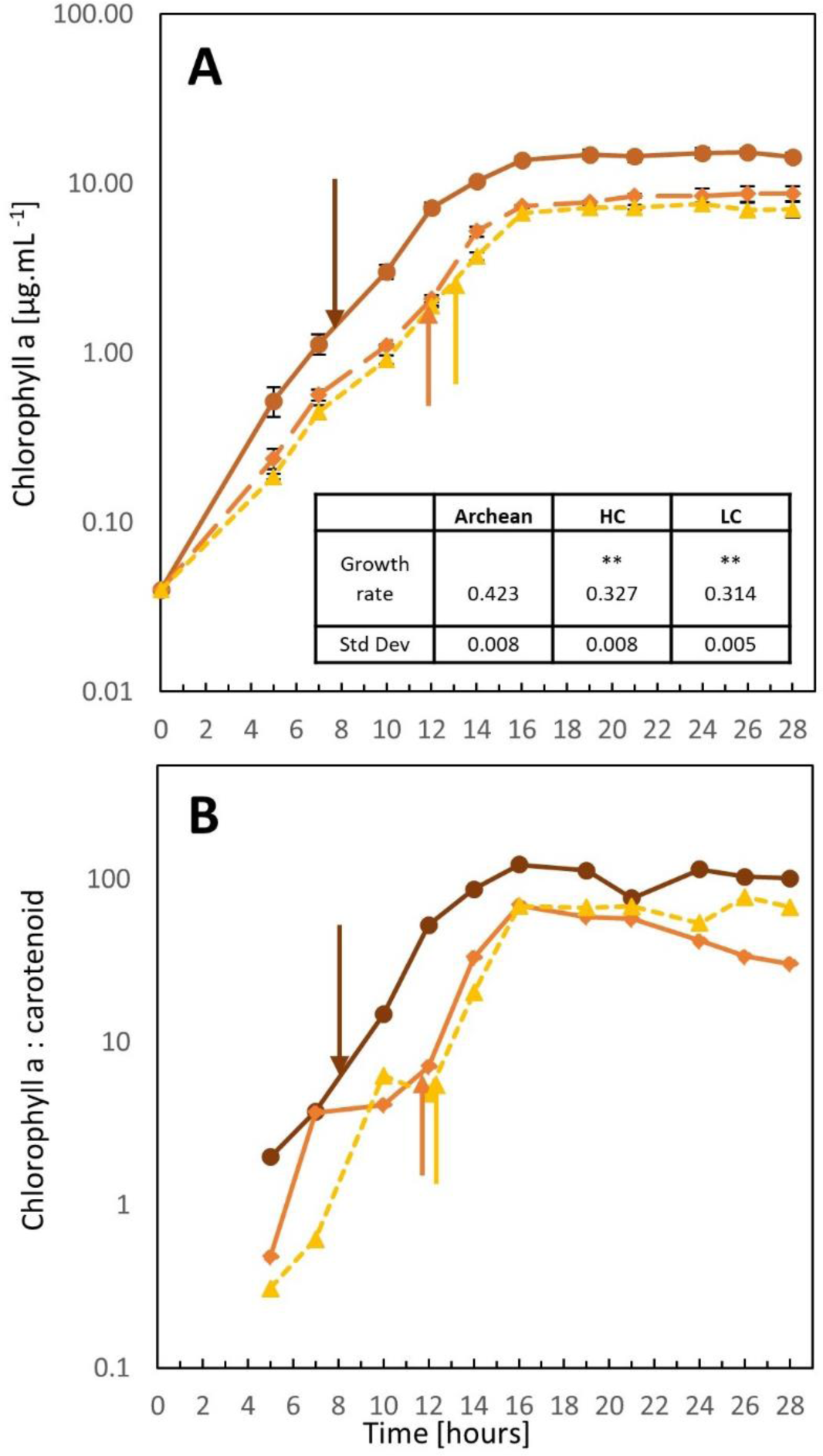
Growth assessment of *Pseudanabaena* sp. PCC7367 under three atmospheric conditions. Triplicate cultures of *Pseudanabena* sp. PCC7367 were inoculated at 0.04 µg. ml^-1^ of Chlorophyll a and monitored for 28 days for chlorophyll a content (A) and used to calculate the growth rate (inset table) for days 0-12. The Chl a: carotenoid content ratios are presented (B). Samples were collected at similar Chl a content from unstirred cultures for RNA extraction and enzyme activity determinations on day 8-9 under Archean atmospheric conditions 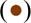, day 12-13 for the high CO_2_ atmosphere grown cultures 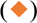 and day 13-14 for normal atmospheric conditions 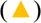, respectively indicated by arrows. Bars represent the standard deviation (n=3). ** p<0.001 for increased growth rate under Archean conditions compared to HC and LC (Pairwise multiple comparison, Holm-Sidak method; Supplementary. Table 6).

### Medium oxygen level assessments

Given that O_2_ accumulation both within and outside the cell, influences SOD expression levels (Kim & Suh, 2005), and as *Pseudanabaena* PCC7367 releases significantly more O_2_ under anoxic conditions (Herrmann et al., 2021), the residual O_2_ levels in the medium was tracked over a full day cycle. In order to obtain basal levels of O_2_ retention in agitated cultures, the O_2_ levels in gently stirred cultures was measured for an additional 24 hrs. Oxygen levels in the medium were consistently higher during the period of active photosynthesis, than during the night, for all conditions measured (Supplementary Fig. 2; Supplementary Table 7). Standard deviations of triplicate measurements were large for stationary cultures, illustrating the influence of O_2_ bubble attachment to the sensors, or clumping of culture mass on or near the sensor. Stirred cultures had less variation between triplicate measurements, reflected in smaller standard deviations (Supplementary Fig. 7). Medium O_2_ levels revealed no significant differences between cultures grown under anoxic or oxygen rich conditions during the day (Figure 3) at time points 2 and 14 hrs. A significant reduction in medium O_2_ levels was recorded for stationary cultures grown under the Archean anoxic simulation compared to LC culture conditions during the night time cycle at 18 (p=0.012) and 22 (p=0.04) hrs (Supplementary table 7). Moreover, stirred cultures of Archean anoxic simulation were significantly reduced in O_2_ levels at night under anoxic conditions, compared to LC (p<0.001) and HC (p=0.001) grown cultures (Supplementary Table 7).

**Figure 3:**
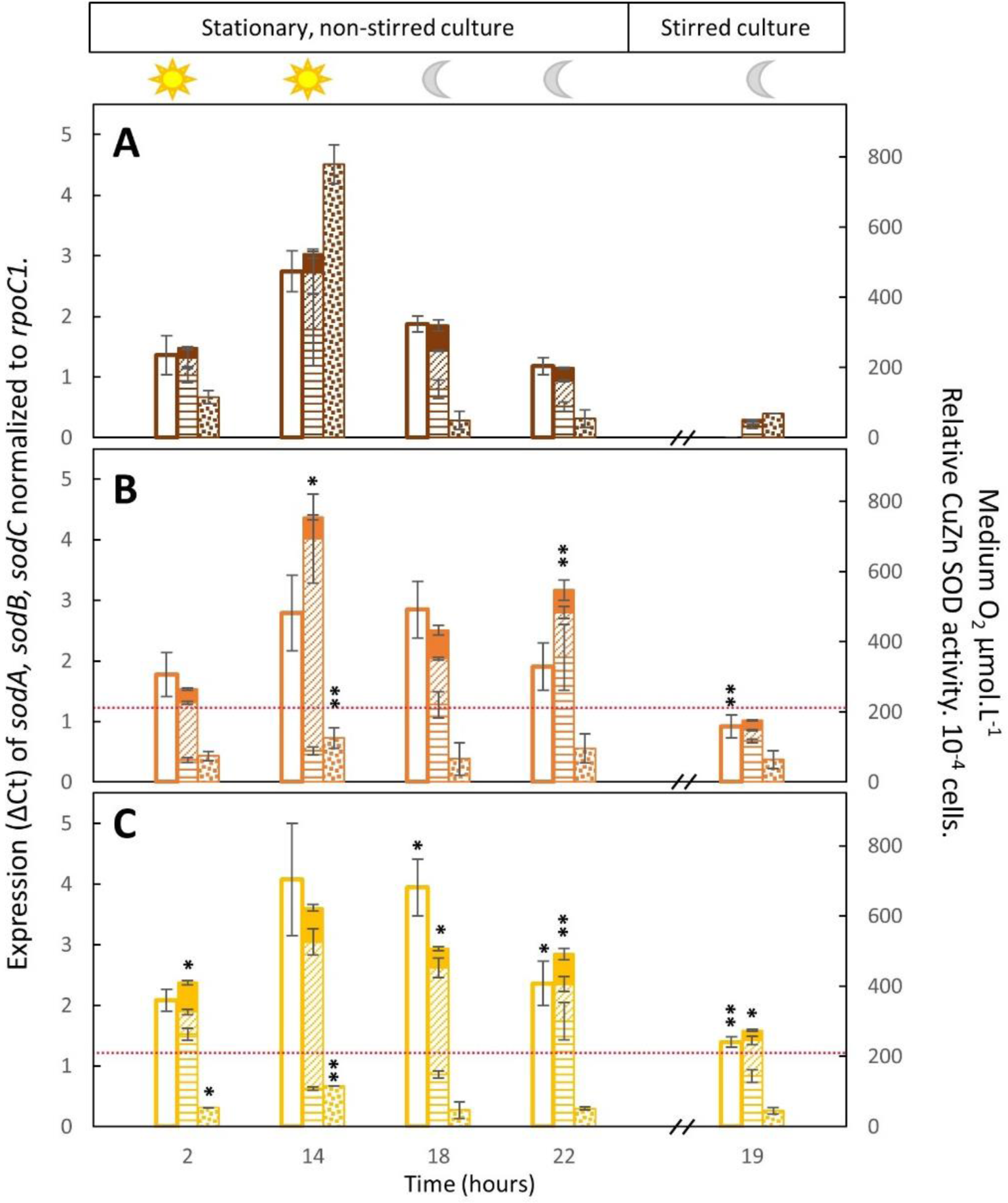
Expression and activity of cytosolic superoxide dismutase against medium O_2_ concentration. Expression of the individual genes, namely *sodA*, encoding a putative MnSOD 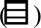, *sodB*, encoding FeSOD 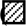 and *sodC*, encoding a putative CuZnSOD 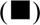 is plotted with the concentration of O_2_ recorded in the medium 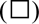 and the relative cytosolic SOD activity 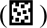 for conditions simulating the anoxic Archean atmosphere (A) and present day oxic conditions with (B) or without (C) CO_2_ supplementation. Data is presented for the first 4 sampling timepoints, namely two hrs after the lights went on (2 hrs), two hrs before they went off (14), two hrs after they went off (18) and 2 hrs before the lights went on again (22 hrs). The cultures were subsequently stirred, and the last sample obtained after 19 hrs, in the dark, to obtain a reading equilibrated with the atmosphere, not influenced by photosynthesis. Bars represent the average of three biological replicates and their standard deviation. The dotted red line indicates the dissolved O_2_ levels of 206 µmol.L^-1^ in artificial seawater medium. The O_2_ concentration was below the detection limit in the Archean simulation experiments at sample point 19. Significant differences of LC and HC to Archean parameters are provided with * for p< 0.05 and ** for p≤ 0.001 (Suppl. Table 8).

*sodABC* Gene Expression and activity assessment in *Pseudanabaena* sp. PCC7367.

To determine whether atmospheric oxygen levels influenced transcription of *sodABC* genes, their expression levels were assessed over 24 hrs. The genome of *Pseudanabaena* sp. PCC7367 (NC_019701.1) includes three SODs, namely a 696 bp *sodC* (Pse7367_0398) encoding a CuZnSOD, a *sodB* (PSE7367_RS14055) of 600 bp for FeSOD and a 765 bp gene, *sodA* (Pse7367_0596), for MnSOD (Boden et al., 2021; Harada et al., 2021). Primers to each gene were validated (Supplementary Tables 1 and 2) and used to quantify SOD gene expression over 24 hrs in non-stirred cultures of *Pseudanabaena* sp. PCC7367 grown under the three atmospheres of the anoxic Archean and oxic LC and HC conditions. Additional samples were collected of stirred cultures, at night, representing the lowest levels of O_2_ in the medium, and hence the potentially lowest O_2_^•-^ levels (Supplementary Fig. 2).

Total expression of all three genes, *sodABC,* in *Pseudanabaena* sp. PCC 7367 correlates significantly (≙SOD sum, R^2^= 0.751, p= 2.81 x 10^-9^; Supplementary Table 9) with the dissolved O_2_ levels in the medium, with the highest expression recorded at 14 hrs (Figure 3), two hrs before the lights went off, under all three atmospheric conditions investigated. The highest expression of *sodABC* in relation to the housekeeping gene, *rpoC1*, was observed for *Pseudanabaena* sp. PCC7367 grown under HC, and was significantly more (p = 0.002; Supplementary Table 8), than under Archean conditions, after 14 hrs of light (Fig. 3A). Transcription of *sodABC* remained significantly raised at LC conditions at night at time point 18 hrs (p=0.014) and 22 hrs (p<0.001) and HC at 22 hrs (p< 0.001) when compared to night time transcription levels under the Archean simulated atmosphere.

Expression of the individual *sodB* and *sodC* genes was highly significantly correlated (Supplementary Table 9) to medium oxygen levels (R^2^= 0.651, p = 2.7 x 10^-7^ and R^2^= 0.675, p = 3.76 x 10^-7^ respectively), with expression of *sodC* correlating strongly to expression of *sodA* (p<0.001) and *sodB* (p<0.001). While *sodA* showed no significant correlation of expression to medium oxygen levels (p= 0.122), it was the only SOD gene that correlated significantly (p=0.019) to total cytosolic SOD activity (Supplementary Table 9). The expression of *sodA* was also significantly higher under Archean conditions at 2 hrs, compared to the HC (p<0.001) and LC (p<0.001) conditions, with LC expression levels also significantly raised compared to HC conditions (p<0.001). Expression of *sodA* was also significantly raised at 14 hr (p=0.02), compared to HC conditions, and afterwards significantly reduced at 18 hr (p=0.03), 22 hr (p=0.006) and 19 hr (p<0.001), as well as LC conditions at 22 hr (p=0.012) and 19 hr (p<0.001).

In contrast to *sodABC* expression levels, a significant (p< 0.001) reduction in cytosolic SOD activity was recorded under HC and LC conditions at 14 hrs, and at 2 hrs (LC only), compared to Archean culture conditions, suggesting impaired enzyme activity high atmospheric O_2_ conditions. While no significant change in cytosolic SOD activity was observed for samples taken after 19 hrs agitation between all three atmospheres, expression of *sodABC* was significantly raised (p=0.03) under LC stirred conditions (Fig. 2C) compared to the anoxic (Fig. 2A) Archean simulation, reflecting the significantly raised levels of dissolved O_2_ in the medium. The lowest transcription of *sodABC* was measured in stirred cultures grown anoxically, three hrs after the lights went out (Fig. 3A) when no free O_2_ was recorded in the medium.

In summary, expression of *sodABC* under Archean conditions, as well as expression of individual *sodB* and *sodC* genes, is significantly reduced (p<0.001) when compared to the modern day oxygen rich atmosphere. Expression of individual *sodB* and *sodC* genes correlate significantly to medium O_2_ levels, whereas expression of *sodA* does not. Additionally, cytosolic SOD activity correlates significantly (p=0.019) to expression of the MnSOD, *sodA*, but neither to *sodB,* nor *sodC* expression.

### Protein properties and modelling

Given the discrepancies in transcriptional levels of the *sodABC* genes in *Pseudanabaena* sp. PCC7367, and their cytosolic activities, analysis of the signal peptide targeting sequences of the SOD proteins was undertaken. The goal was to ascertain whether SODs could be exported from the cytoplasm to be inserted in the thylakoid or cell membranes or targeted to the thylakoid lumen or periplasmic space. H_2_O_2_ can be actively or passively transported across cellular membranes, but O_2_^•-^ is unable to cross lipid membranes (Hansel & Diaz, 2021). Targeting a SOD to the lumen of the thylakoid, to the site of O_2_^•-^ generation, may offer protection from superoxide induced damage.

MnSOD of *Pseudanabaena* sp. PCC7367 has a predicted signal peptide sequence containing a twin arginine motif (RR), followed by a hydrophobic segment and a A-x-A peptidase cleavage site (Fig. 4A). This indicates that MnSOD may be a substrate of the TAT protein translocation system that transports proteins in their folded state into either the periplasmic space, or the thylakoid lumen, or both. Homologues to the *E. coli* Tat system were also identified in *Pseudanabaena* sp. PCC7367, namely two Tat A homologues and a single Tat C homologue, suggesting that it has a minimal Tat translocation system (Russo & Zedler, 2021). The MnSOD protein sequence also carries a second methionine right after the predicted peptidase cleavage site, which may be an alternative start codon for a cytosolic form of MnSOD, starting with the sequence MGLT.

**Figure 4.**
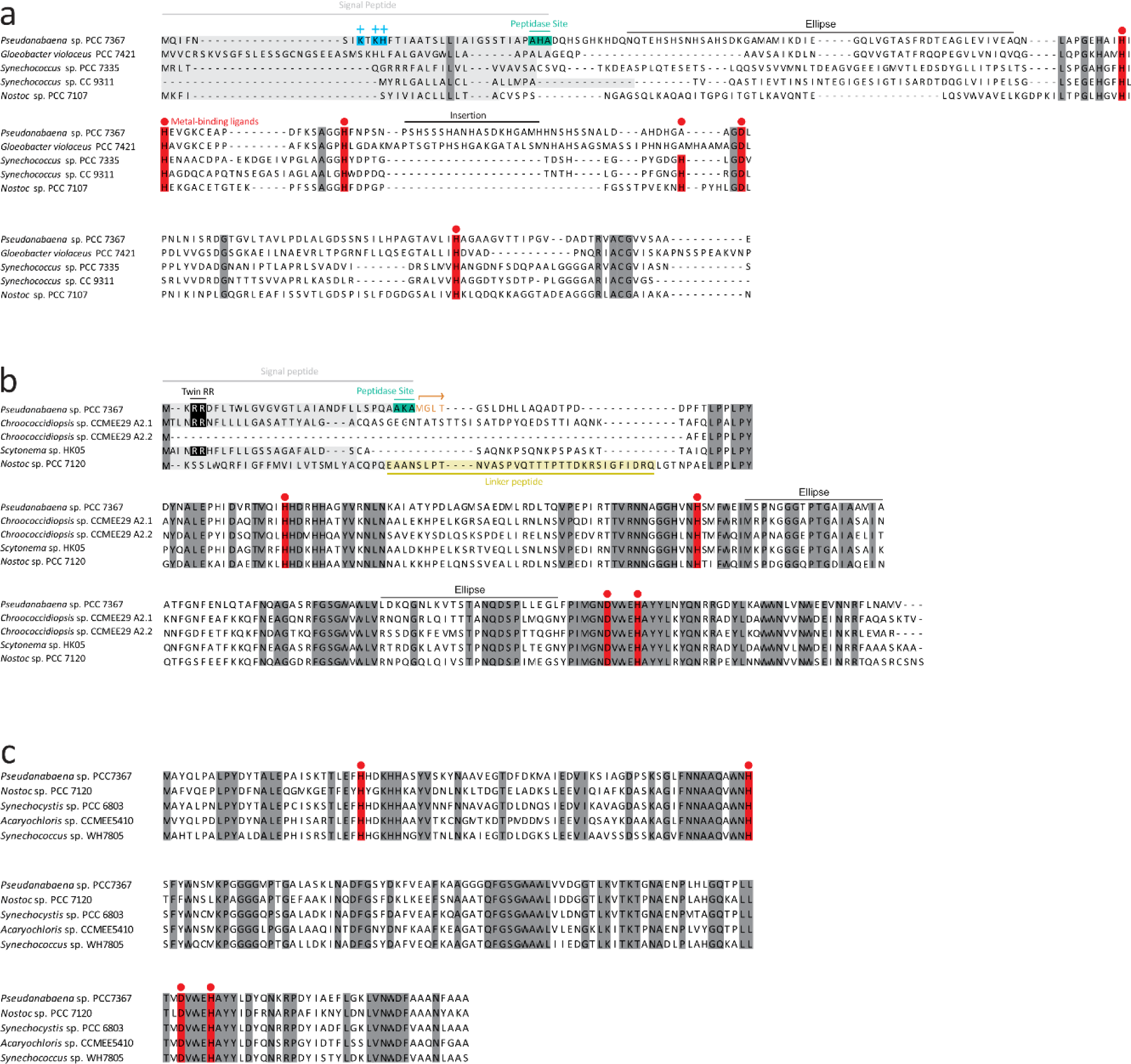
Multiple sequence alignments of the super oxide dismutases encoded on the genome of *Pseudanabaena* sp. PCC7367. a) The CuZnSOD protein sequence has a positively charged N-terminal sequence followed by a potential transmembrane segment and AxA signal peptidase cleavage site, which suggests export from the cytosol via the Sec secretory system. b) The N-terminus of the *Pseudanabaena* MnSOD carries a potential twin arginine signal (twin RR), transmembrane segment, and signal peptidase cleavage site (<u>AxA</u>). The second methionine residue following the cleavage site could indicate a second site of initiation of translation, which may produce a cytosolic form of MnSOD starting with MGLT (highlighted in orange). c) The FeSOD lacks a potential targeting sequence. Metal-binding sites are highlighted in red. NCBI accessions of each sequence are as follows: CuZnSODS (WP_015163683.1, BAC89922.1, WP_006457095.1, WP_011619688.1 and WP_015114498.1), MnSODs (AFY68900.1, QUX80117.1, WP_250121708.1, WP_073632316.1 and WP_010994247.1), FeSODS (WP_015166022.1, WP_010997089.1, WP_010872652.1, WP_010469685.1 and WP_006041934.1).

CuZnSOD has a positively charged N-terminal sequence followed by a hydrophobic segment (Fig. 4B), a classical signal peptide for the Sec translocation system. This suggests that CuZnSOD may be transported through the plasma membrane and/or the thylakoid membrane of *Pseudanabaena* sp. PCC7367 and hence be in the periplasmic space and/or the thylakoid lumen. Alignment of related protein sequences for cyanobacterial SodC, highlighted a 23 amino acid insertion sequence, that may interfere with the binding of Zn^2+^ in the enzyme reaction centre, thereby potentially reducing its activity (Fig. 5).

**Figure 5.**
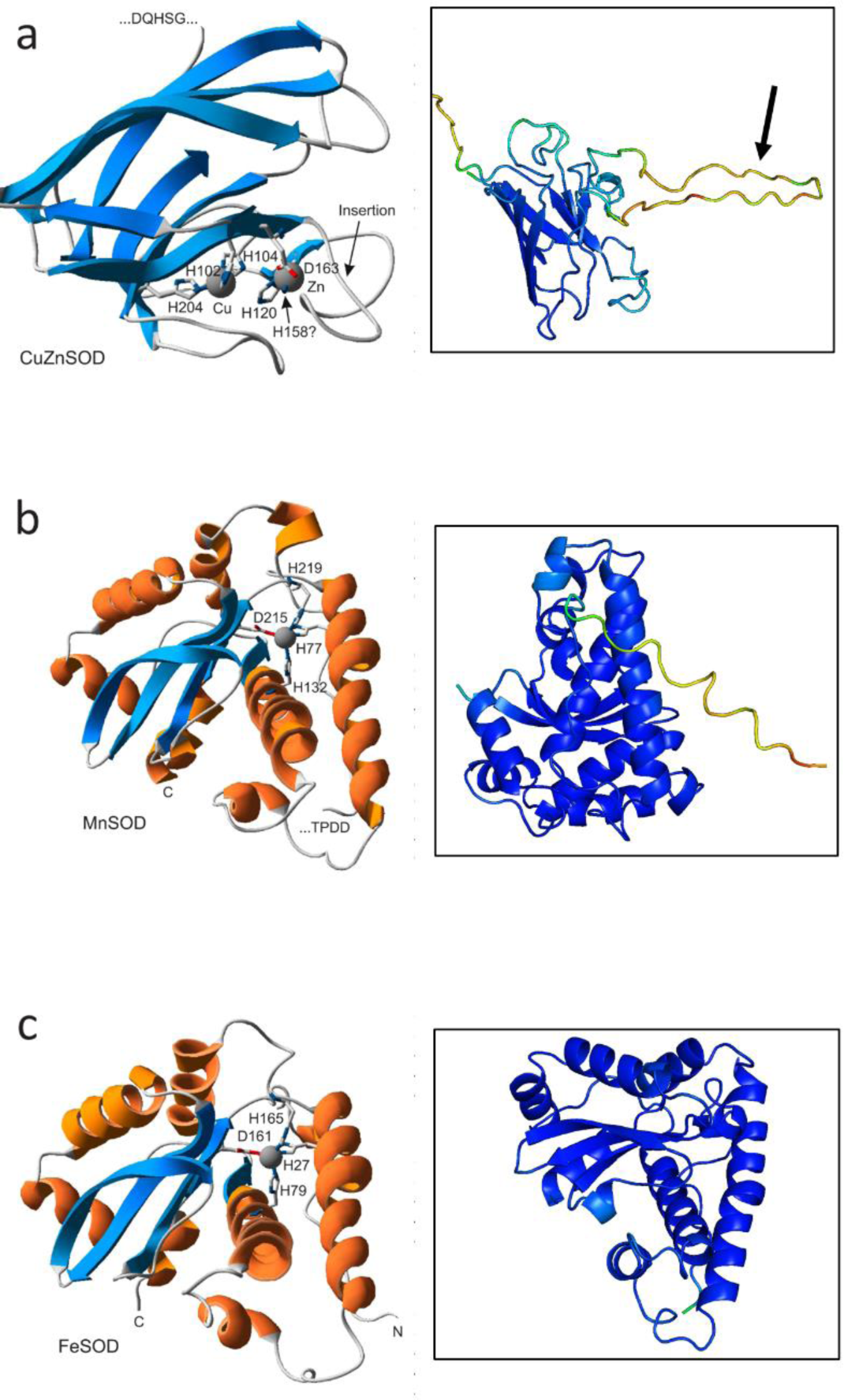
Predicted protein structures of super oxide dismutases encoded on the genome of *Pseudanabaena* sp. PCC7367. a). The CuZnSOD protein prediction of the truncated protein cleaved at the AxA signal peptidase cleavage site. The arrows indicate the 23 aa insertion. The N-ternimal is cutof in this frame. b) *Pseudanabaena* MnSOD cleaved at the signal peptidase cleavage site (<u>AxA</u>). c) The structural predictions for the complete protein sequence for FeSOD. The models on the left were generated at the Phyre2 model server (L. A. Kelley et al., 2015) and visualized using the Swiss-PDB viewer (Guex et al., 2009). Figures on the right were generated using Alphafold2 (Jumper et al., 2021; Varadi et al., 2021) and shaded using PYMOL2 (DeLano, 2020). Blue shading indicates high confidence in structure (100 %) with decreasing confidence through the spectrum to deep red, indicating a confidence of 25%.

FeSOD was found to have no positively charged N-terminus and no hydrophobic signal, suggesting it is a soluble cytosolic protein (Fig. 4C).

The predicted protein structures using Alphafold 2 (Fig. 5) indicated high confidence in the structures for the FeSOD and the truncated MnSOD sequence generated, to existing functional enzymes (Supplementary Fig. 6 for multiple sequence alignments), suggesting they are indeed active in the cytoplasm and thylakoid/periplasmic space respectively. The 23 aa insertion observed in the CuZnSOD sequence is not a common feature, however the predicted structure of the truncated core enzyme has high congruence to existing CuZnSODs (Fig. 5a). The insertion may, however, interfere with the binding of Zn in the active site, thereby possibly reducing its activity.

Furthermore, screening of the *Pseudanabaena* sp. PCC7367 genome indicated a complete complement of metal ion transporters to support the synthesis of MnSOD and CuZnSOD (Supplementary Table 4). The iron transporters were previously identified in *Pseudanabaena* sp. PCC7367 (Enzingmüller-Bleyl et al., 2022). Additionally, no genes encoding a SOR were found using tBLASTn, nor were they annotated as such in KEGG, so all superoxide inactivation must be a result of the SODs presented above. No homologue to the *E. coli* catalase gene was identified (Harada et al., 2021). However, the genome of *Pseudanabaena* sp. PCC7367 encodes five predicted peroxiredoxins, a presumably thioredoxin-dependent glutathione peroxidase and one predicted heme-containing peroxidase that could reduce the SOD dismutation product H_2_O_2_ (Supplementary Table 5).

## Discussion

Life on early Earth, prior to the oxygenation of the atmosphere during the GOE, was different to that found today (Fischer et al., 2016; Sessions et al., 2009). Cyanobacteria, the only known prokaryotes capable of conducting oxygenic photosynthesis are considered the primary agents for bringing about this change through the release of free O_2_ to the atmosphere by the photocatalytic hydrolysis of water (Cardona et al., 2019). The increase in levels of free oxygen, and its accompanying superoxide radical, raised the question as to whether early Cyanobacteria were able to inactivate O_2_^•-^ arising from oxygenic photosynthesis ref (Fischer & Valentine, 2019; Hamilton, 2019). The phylogenetic trajectory of CuZnSODs matches that of the deep branching Cyanobacteria at the base of the genomic tree, suggesting that these early strains were able to process O_2_^•-^ prior to the GOE (Boden et al., 2021; Harada et al., 2021; Ślesak et al., 2019). Elevated levels of O_2_ per Chl a were recorded for a deep branching, CuZnSOD encoding Cyanobacterium, *Pseudanabaena* sp. PCC7367 under an Archean simulated, anoxic atmosphere (Herrmann et al., 2021). Hence, in this study we explored whether an increase in oxygen production is accompanied with increased levels of expression and activities of SODs, specifically CuZnSOD.

Unexpectedly, this study revealed a significantly increased growth rate for *Pseudanabaena* sp. PCC7367 cultured under an anoxic atmosphere, accompanied by significant increases in both viability markers of glycogen and protein. Additionally, there was no significant difference in accumulation of dissolved oxygen in the medium of stationary cultures during the day between all three atmospheres tested, however, significant differences were recorded at night between the Archean simulation and current LC conditions. Agitation of the cultures simulated a natural, shallow-water oxygen oases described in the literature (Crowe et al., 2013; Eickmann et al., 2018; Herrmann et al., 2021; Homann et al., 2015), suggesting that early cyanobacteria would not have been subjected to high O_2_ levels at night, allowing them time to recover from daytime photosynthetically generated free radicals.

Cyanobacterial Chl *a* content is commonly used to assess filamentous cyanobacterial growth (Li et al., 2014), while the carotenoid content provides an indication of potential light stress in cyanobacteria capable of synthesising this pigment (Herrmann & Gehringer, 2019). Additionally, carotenoids dissipate excess light energy, thereby scavenging O_2_^•-^ radicals in the cytoplasm (Latifi et al., 2009). As no significant change in the carotenoid content was observed, we can assume that carotenoid production was not upregulated to compensate for additional free radical removal.

Furthermore, this study demonstrates enhanced total expression of *sodABC* in culture material harvested towards day’s end, corresponding to high levels of O_2_ in the medium under all three atmospheres investigated. This agrees with studies on *Synechocystis* under LC conditions (Kim & Suh, 2005) and literature reporting O_2_^•-^ levels peaking near midday in surface waters in the ocean, and ponds (Zinser, 2018). Specifically, *sodB* and *sodC* expression, encoding the FeSOD and CuZnSOD respectively, correlates significantly to levels of dissolved O_2_ in the medium. The increased SOD activity observed during the day in cultures growing under the Archean simulated anoxic atmosphere corresponds with the increased O_2_ release rates per Chl a content recorded (Herrmann et al., 2021) in cultures of *Pseudanabaena* sp. PCC7367 grown under the same experimental conditions. In contrast, expression of *sodA*, encoding MnSOD, correlated significantly to cytosolic SOD activity.

Transcription of *sodABC* increased in agitated HC and LC cultures at 19 hrs, compared to cultures under anoxic conditions, with no significant difference in cytosolic SOD activity observed. While transcription levels often do not correlate to translation rates or protein activity, due to mRNA stability or translational regulation, our data suggests cyanobacteria are constantly challenged growing under modern-day levels of oxygen. They consequently require inactivation of O_2_^•-^, even at night, when no oxygen is being released from photosynthesis.

As total *sodABC* transcription levels did not correlate to cytosolic SOD activity, we explored whether the encoded SOD isoforms could be exported out of the cytoplasm, to either the thylakoid or periplasmic space, or remain membrane-embedded. The genome of *Pseudanabaena* sp. PCC7367 carries a single homologue of SecA, SecD, SecY, SecE and SecG, with two homologues of SecF (Supplementary Table 3), suggesting it carries a functional Sec export pathway (Russo & Zedler, 2021). Homologues to the *E. coli* Tat system were also identified in *Pseudanabaena* sp. PCC7367, namely two Tat A homologues and a single Tat C homologue, suggesting that it has a minimal Tat translocation system (Russo & Zedler, 2021). Analysis of the three putative SOD isoforms encoded by *Pseudanabaena* sp. PCC7367 suggested that there were potentially two active SODs in the cytoplasm. The FeSOD protein, which carries no signal peptide, would remain in the cytosol. This is in agreement with the cytosolic location of FeSOD in *Synechocystis* (Ke et al., 2014; Kim & Suh, 2005). Synthesis of MnSOD could be initiated from a second translation initiation site to generate a protein that would remain in the cytoplasm. Additionally, the longer protein carries a twin arginine motif, similar to that found in *Chroococcidiopsis* (Napoli et al., 2021), indicating that it may be transported through the thylakoid and / or cell membrane to function in the thylakoid lumen, or periplasmic space, respectively. Multiple forms of MnSOD have been isolated from the thylakoid membrane, lumen or cytoplasm (Herbert et al., 1992; Raghavan et al., 2013; Raghavan et al., 2015). Given that the expression of MnSOD did not correlate to external dissolved O_2_ levels in the media, we propose that the MnSOD is predominantly targeted to the thylakoid lumen to inactivate reactive O_2_^•-^ species at the site where they are generated during photosynthesis. A correlation between the cytosolic SOD activity and the expression of *sodA* might also point to a partial cytosolic localization of MnSOD similar to *Nostoc* sp. PCC7120 (Raghavan et al., 2013; Raghavan et al., 2015).

The CuZnSOD from *Pseudanabaena* sp. PCC7367 has no secondary translation site and, as it carries a Sec translocation signal peptide, would therefore presumably always be transported across the cell and / or thylakoid membrane. In contrast, the *sodC*-encoded protein from *Chroococcidiopsis* does not carry a signal peptide and is assumed to be cytosolic (Napoli et al., 2021). Given that expression of *Pseudanabaena* sp. PCC7367 *sodC* strongly correlated to external medium O_2_ levels, we propose that this SOD isoform is exported across the cell membrane to reduce the levels of O_2_^•-^ in the periplasm.

**In conclusion,** this study demonstrates that the early branching cyanobacterium, *Pseudanabaena* sp. PCC7367, exhibits increased cytoplasmic SOD activity under the anoxic atmospheric conditions that existed on early Earth, prior to the GOE. Additionally, cultures grown under Archean atmospheric conditions grow faster and contain significantly more protein and glycogen, thereby contributing valuable biologically available C & N to the environment compared to cultures grown under modern-day oxic conditions. These data show that aquatic marine Cyanobacteria growing under present day atmospheric levels of O_2_ are exposed to significantly higher levels of dissolved oxygen, thereby potentially placing a higher O_2_^•-^ load on the cellular metabolism, especially at night. Increased transcription of SOD genes without the corresponding increase in cytosolic activity may suggest a higher enzyme turnover rate. In summary, this investigation helps us understand the complex interplay between O_2_ production, O_2_^•-^ and cellular protection mechanisms in early cyanobacteria, offering valuable insights into the evolutionary mechanisms that shaped our planet’s biosphere. Further investigations into the functionality of SOD isoforms will help us understand how these ancient organisms navigated the transition to O_2_-rich environments, laying the foundation for the diverse life forms that thrive in today’s oxygenated atmosphere.

## Acknowledgements

This investigation was funded by the German Research Foundation (DFG) under the SPP1833 grants GE2558/3-1 & GE2558/4-1 awarded to MMG, and a Royal Society University Research Fellowship awarded to P.S-B, a University of Bristol Graduate Teaching Scholarship awarded to J.S.B. with additional funding from a NERC Frontiers grant (NE/V010824/1) awarded to Dr Eva E. Stüeken at the University of St. Andrews. S.S.T. was partly funded by the DFG RTG 2737 (Stressistance).

## Author contributions

Conceptualization-J.S.B, P.S-B & M.M.G; Investigation-S.S.T., J.S.B, M.D., K.M.K. & J.H; Formal analysis-S.T., J.S.B., K.M.K., N.W., J.H., M.D. & M.M.G.; Resources - N.F-D & M.M.G; and Writing - review & editing - all authors.

## References

Alexova, R., Fujii, M., Birch, D., Cheng, J., Waite, T. D., Ferrari, B. C., & Neilan, B. A. (2011). Iron uptake and toxin synthesis in the bloom-forming Microcystis aeruginosa under iron limitation. Environmental Microbiology, 13(4), 1064–1077.

Altschul, S. F. (1991). Amino acid substitution matrices from an information theoretic perspective. Journal of molecular biology, 219(3), 555–565.

Bartsevich, V. V., & Pakrasi, H. (1995). Molecular identification of an ABC transporter complex for manganese: analysis of a cyanobacterial mutant strain impaired in the photosynthetic oxygen evolution process. The EMBO journal, 14(9), 1845–1853.

Bekker, A., Holland, H. D., Wang, P. L., Rumble, D., Stein, H. J., Hannah, J. L., Coetzee, L. L., & Beukes, N. J. (2004). Dating the rise of atmospheric oxygen. Nature, 427(6970), 117–120. 10.1038/nature02260

Boden, J. S., Konhauser, K. O., Robbins, L. J., & Sánchez-Baracaldo, P. (2021). Timing the evolution of antioxidant enzymes in cyanobacteria [Article]. Nature Communications, 12(1), Article 4742. 10.1038/s41467-021-24396-y

Cardona, T., Sánchez-Baracaldo, P., Rutherford, A. W., & Larkum, A. W. (2019). Early Archean origin of photosystem II. Geobiology, 17(2), 127–150.

Case, A. J. (2017). On the origin of superoxide dismutase: an evolutionary perspective of superoxide-mediated redox signaling. Antioxidants, 6(4), 82.

Catling, D. C., & Zahnle, K. J. (2020). The archean atmosphere. Science Advances, 6(9), eaax1420.

Crowe, S. A., Døssing, L. N., Beukes, N. J., Bau, M., Kruger, S. J., Frei, R., & Canfield, D. E. (2013). Atmospheric oxygenation three billion years ago. Nature, 501(7468), 535–538. 10.1038/nature12426

DeLano, W. L. (2020). PyMOL Molecular Graphics System. In Schrödinger LLC.

Dupont, C., Neupane, K., Shearer, J., & Palenik, B. (2008). Diversity, function and evolution of genes coding for putative Ni-containing superoxide dismutases. Environmental Microbiology, 10(7), 1831–1843.

Eickmann, B., Hofmann, A., Wille, M., Bui, T. H., Wing, B. A., & Schoenberg, R. (2018). Isotopic evidence for oxygenated Mesoarchaean shallow oceans. Nature Geoscience, 11(2), 133–138. 10.1038/s41561-017-0036-x

Enzingmüller-Bleyl, T. C., Boden, J. S., Herrmann, A. J., Ebel, K. W., Sánchez-Baracaldo, P., Frankenberg-Dinkel, N., & Gehringer, M. M. (2022). On the trail of iron uptake in ancestral Cyanobacteria on early Earth. Geobiology.

Fischer, W. W., Hemp, J., & Valentine, J. S. (2016). How did life survive Earth’s great oxygenation? Current opinion in chemical biology, 31, 166–178.

Fischer, W. W., & Valentine, J. S. (2019). How did life come to tolerate and thrive in an oxygenated world? Free Radical Biology and Medicine, 140, 1–3. 10.1016/j.freeradbiomed.2019.07.021

Fournier, G., Moore, K., Rangel, L., Payette, J., Momper, L., & Bosak, T. (2021). The Archean origin of oxygenic photosynthesis and extant cyanobacterial lineages. Proceedings of the Royal Society B, 288(1959), 20210675.

Guex, N., Peitsch, M. C., & Schwede, T. (2009). Automated comparative protein structure modeling with SWISS-MODEL and Swiss-PdbViewer: A historical perspective. ELECTROPHORESIS, 30(S1), S162–S173. 10.1002/elps.200900140

Hallgren, J., Tsirigos, K. D., Pedersen, M. D., Armenteros, J. J. A., Marcatili, P., Nielsen, H., Krogh, A., & Winther, O. (2022). DeepTMHMM predicts alpha and beta transmembrane proteins using deep neural networks. bioRxiv, 2022.2004.2008.487609. 10.1101/2022.04.08.487609

Hamilton, T. L. (2019). The trouble with oxygen: The ecophysiology of extant phototrophs and implications for the evolution of oxygenic photosynthesis. Free Radical Biology and Medicine, 140, 233–249. 10.1016/j.freeradbiomed.2019.05.003

Hansel, C. M., & Diaz, J. M. (2021). Production of Extracellular Reactive Oxygen Species by Marine Biota. Annual Review of Marine Science, 13(1), 177–200. 10.1146/annurev-marine-041320-102550

Harada, M., Akiyama, A., Furukawa, R., Yokobori, S. I., Tajika, E., & Yamagishi, A. (2021). Evolution of Superoxide Dismutases and Catalases in Cyanobacteria: Occurrence of the Antioxidant Enzyme Genes before the Rise of Atmospheric Oxygen [Article]. Journal of Molecular Evolution, 89(8), 527–543. 10.1007/s00239-021-10021-5

Herbert, S. K., Samson, G., Fork, D. C., & Laudenbach, D. E. (1992). Characterization of damage to photosystems I and II in a cyanobacterium lacking detectable iron superoxide dismutase activity. Proceedings of the National Academy of Sciences, 89(18), 8716–8720.

Herrmann, A., Sorwat, J., Byrne, J., Frankenberg-Dinkel, N., & Gehringer, M. (2021). Diurnal Fe (II)/Fe (III) cycling and enhanced O2 production in a simulated Archean marine oxygen oasis. Nature Communications, 12(1), 1–11.

Herrmann, A. J., & Gehringer, M. M. (2019). An investigation into the effects of increasing salinity on photosynthesis in freshwater unicellular cyanobacteria during the late Archaean. Geobiology, 17(4), 343–359.

Homann, M., Heubeck, C., Airo, A., & Tice, M. M. (2015). Morphological adaptations of 3.22 Ga-old tufted microbial mats to Archean coastal habitats (Moodies Group, Barberton Greenstone Belt, South Africa). Precambrian Research, 266, 47–64. 10.1016/j.precamres.2015.04.018

Ismaiel, M., El-Ayouty, Y. M., Loewen, P. C., & Piercey-Normore, M. D. (2014). Characterization of the iron-containing superoxide dismutase and its response to stress in cyanobacterium Spirulina (Arthrospira) platensis. Journal of Applied Phycology, 26(4), 1649–1658.

Jabłońska, J., & Tawfik, D. S. (2021). The evolution of oxygen-utilizing enzymes suggests early biosphere oxygenation. Nature Ecology & Evolution, 5(4), 442–448. 10.1038/s41559-020-01386-9

Jahodářová, E., Dvořák, P., Hašler, P., Holušová, K., & Poulíčková, A. (2018). Elainella gen. nov.: a new tropical cyanobacterium characterized using a complex genomic approach. European Journal of Phycology, 53(1), 39–51.

Johnson, L. A., & Hug, L. A. (2019). Distribution of reactive oxygen species defense mechanisms across domain bacteria. Free Radical Biology and Medicine, 140, 93–102. 10.1016/j.freeradbiomed.2019.03.032

Jumper, J., Evans, R., Pritzel, A., Green, T., Figurnov, M., Ronneberger, O., Tunyasuvunakool, K., Bates, R., Žídek, A., & Potapenko, A. (2021). Highly accurate protein structure prediction with AlphaFold. Nature, 596(7873), 583–589.

Kanehisa, M., Furumichi, M., Sato, Y., Kawashima, M., & Ishiguro-Watanabe, M. (2022). KEGG for taxonomy-based analysis of pathways and genomes. Nucleic acids research, 51(D1), D587–D592. 10.1093/nar/gkac963

Ke, W.-T., Dai, G.-Z., Jiang, H.-B., Zhang, R., & Qiu, B.-S. (2014). Essential roles of iron superoxide dismutase in photoautotrophic growth of Synechocystis sp. PCC 6803 and heterogeneous expression of marine Synechococcus sp. CC9311 copper/zinc superoxide dismutase within its sodB knockdown mutant. Microbiology, 160(11), 228-241.

Kelley, L. A., Mezulis, S., Yates, C. M., Wass, M. N., & Sternberg, M. J. (2015). The Phyre2 web portal for protein modeling, prediction and analysis. Nat Protoc, 10(6), 845–858. 10.1038/nprot.2015.053

Kelley, L. A., Mezulis, S., Yates, C. M., Wass, M. N., & Sternberg, M. J. E. (2015). The Phyre2 web portal for protein modeling, prediction and analysis. Nature Protocols, 10(6), 845–858. 10.1038/nprot.2015.053

Kim, J.-H., & Suh, K. H. (2005). Light-dependent expression of superoxide dismutase from cyanobacterium Synechocystis sp. strain PCC 6803. Archives of Microbiology, 183(3), 218–223.

Klotz, A., Georg, J., Bučinská, L., Watanabe, S., Reimann, V., Januszewski, W., Sobotka, R., Jendrossek, D., Hess, W. R., & Forchhammer, K. (2016). Awakening of a dormant cyanobacterium from nitrogen chlorosis reveals a genetically determined program. Current biology, 26(21), 2862–2872.

Kump, L. R. (2008). The rise of atmospheric oxygen. Nature, 451(7176), 277–278. 10.1038/nature06587

Latifi, A., Ruiz, M., & Zhang, C.-C. (2009). Oxidative stress in cyanobacteria. FEMS microbiology reviews, 33(2), 258–278.

Li, T., Huang, X., Zhou, R., Liu, Y., Li, B., Nomura, C., & Zhao, J. (2002). Differential expression and localization of Mn and Fe superoxide dismutases in the heterocystous cyanobacterium Anabaena sp. strain PCC 7120. Journal of Bacteriology, 184(18), 5096–5103.

Li, Y., Lin, Y., Loughlin, P. C., & Chen, M. (2014). Optimization and effects of different culture conditions on growth of Halomicronema hongdechloris–a filamentous cyanobacterium containing chlorophyll f. Frontiers in plant science, 5, 67.

Lucchetti-Miganeh, C., Goudenège, D., Thybert, D., Salbert, G., & Barloy-Hubler, F. (2011). SORGOdb: superoxide reductase gene ontology curated database. BMC microbiology, 11(1), 1–12.

Lyons, T. W., Reinhard, C. T., & Planavsky, N. J. (2014). The rise of oxygen in Earth’s early ocean and atmosphere. Nature, 506(7488), 307–315.

Meeks, J. C., & Castenholz, R. W. (1971). Growth and photosynthesis in an extreme thermophile, Synechococcus lividus (Cyanophyta). Archiv für Mikrobiologie, 78(1), 25–41.

Miller, A.-F. (2012). Superoxide dismutases: Ancient enzymes and new insights. FEBS letters, 586(5), 585–595. 10.1016/j.febslet.2011.10.048

Napoli, A., Iacovelli, F., Fagliarone, C., Pascarella, G., Falconi, M., & Billi, D. (2021). Genome-Wide Identification and Bioinformatics Characterization of Superoxide Dismutases in the Desiccation-Tolerant Cyanobacterium Chroococcidiopsis sp. CCMEE 029. Frontiers in Microbiology, 12, 660050.

Ouzounis, C. A., Kunin, V., Darzentas, N., & Goldovsky, L. (2006). A minimal estimate for the gene content of the last universal common ancestor—exobiology from a terrestrial perspective. Research in microbiology, 157(1), 57–68.

Peskin, A. V., & Winterbourn, C. C. (2017). Assay of superoxide dismutase activity in a plate assay using WST-1. Free Radical Biology and Medicine, 103, 188–191.

Priya, B., Premanandh, J., Dhanalakshmi, R. T., Seethalakshmi, T., Uma, L., Prabaharan, D., & Subramanian, G. (2007). Comparative analysis of cyanobacterial superoxide dismutases to discriminate canonical forms. BMC Genomics, 8(1), 1–10.

Raghavan, P. S., Rajaram, H., & Apte, S. K. (2013). N-terminal processing of membrane-targeted Mn SOD and formation of multiple active superoxide dismutase dimers in the nitrogen-fixing cyanobacterium Anabaena sp. strain PCC 7120. The FEBS Journal, 280(19), 4827–4838.

Raghavan, P. S., Rajaram, H., & Apte, S. K. (2015). Membrane targeting of MnSOD is essential for oxidative stress tolerance of nitrogen-fixing cultures of Anabaena sp. strain PCC7120. Plant molecular biology, 88(4), 503–514.

Riding, R., Fralick, P., & Liang, L. (2014). Identification of an Archean marine oxygen oasis. Precambrian Research, 251, 232–237.

Russo, D. A., & Zedler, J. A. (2021). Genomic insights into cyanobacterial protein translocation systems. Biological chemistry, 402(1), 39–54.

Sánchez-Baracaldo, P. (2015). Origin of marine planktonic cyanobacteria. Scientific reports, 5(1), 1–10.

Sánchez-Baracaldo, P., Raven, J. A., Pisani, D., & Knoll, A. H. (2017). Early photosynthetic eukaryotes inhabited low-salinity habitats. Proceedings of the National Academy of Sciences, 114(37), E7737–E7745.

Sessions, A. L., Doughty, D. M., Welander, P. V., Summons, R. E., & Newman, D. K. (2009). The continuing puzzle of the great oxidation event. Current biology, 19(14), R567–R574.

Sharon, G., Garg, N., Debelius, J., Knight, R., Dorrestein, P. C., & Mazmanian, S. K. (2014). Specialized metabolites from the microbiome in health and disease. Cell metabolism, 20(5), 719–730.

Shih, P. M., Wu, D., Latifi, A., Axen, S. D., Fewer, D. P., Talla, E., Calteau, A., Cai, F., Tandeau de Marsac, N., & Rippka, R. (2013). Improving the coverage of the cyanobacterial phylum using diversity-driven genome sequencing. Proceedings of the National Academy of Sciences, 110(3), 1053–1058.

Ślesak, I., Kula, M., Ślesak, H., Miszalski, Z., & Strzałka, K. (2019). How to define obligatory anaerobiosis? An evolutionary view on the antioxidant response system and the early stages of the evolution of life on Earth. Free Radical Biology and Medicine, 140, 61–73. 10.1016/j.freeradbiomed.2019.03.004

Ślesak, I., Ślesak, H., Zimak-Piekarczyk, P., & Rozpądek, P. (2016). Enzymatic Antioxidant Systems in Early Anaerobes: Theoretical Considerations [Article]. Astrobiology, 16(5), 348–358. 10.1089/ast.2015.1328

Varadi, M., Anyango, S., Deshpande, M., Nair, S., Natassia, C., Yordanova, G., Yuan, D., Stroe, O., Wood, G., Laydon, A., Žídek, A., Green, T., Tunyasuvunakool, K., Petersen, S., Jumper, J., Clancy, E., Green, R., Vora, A., Lutfi, M.,…Velankar, S. (2021). AlphaFold Protein Structure Database: massively expanding the structural coverage of protein-sequence space with high-accuracy models. Nucleic Acids Research, 50(D1), D439–D444. 10.1093/nar/gkab1061

Ward, L. M., Kirschvink, J. L., & Fischer, W. W. (2016). Timescales of oxygenation following the evolution of oxygenic photosynthesis. Origins of Life and Evolution of Biospheres, 46(1), 51–65.

Warke, M. R., Di Rocco, T., Zerkle, A. L., Lepland, A., Prave, A. R., Martin, A. P., Ueno, Y., Condon, D. J., & Claire, M. W. (2020). The great oxidation event preceded a paleoproterozoic “snowball Earth”. Proceedings of the National Academy of Sciences, 117(24), 13314–13320.

Wellburn, A. R. (1994). The spectral determination of chlorophylls a and b, as well as total carotenoids, using various solvents with spectrophotometers of different resolution. Journal of plant physiology, 144(3), 307–313.

Ye, J., Coulouris, G., Zaretskaya, I., Cutcutache, I., Rozen, S., & Madden, T. L. (2012). Primer-BLAST: a tool to design target-specific primers for polymerase chain reaction. BMC bioinformatics, 13(1), 1–11.

Zinser, E. R. (2018). The microbial contribution to reactive oxygen species dynamics in marine ecosystems. Environmental microbiology reports, 10(4), 412–427.

