## Supplementary figures for "Early-branching cyanobacteria up-regulate superoxide dismutase activity under a simulated early Earth anoxic atmosphere"

**Supplementary Text.**

**Calculation of the relative expression levels of the SOD genes relative to the control housekeeping gene.** The relative expression of the SOD genes relative to *rpoC1* housekeeping gene was calculated as described below. The fold-difference of relative gene expression was calculated from the the threshold cycle number (Ct) which is inversely related to the starting amount of target cDNA (Nolan *et al.*, 2013). The Ct values for the housekeeping gene (HG, *rpoC1*) and the genes of interest (GOI); namely *sodA, sodB* and *sodC* were determined under anoxic Archean simulations, as well as oxic HC and LC conditions. The difference between the HG and the GOI (ΔCt) of the control and test samples was first calculated using the following equation-

$$\Delta Ct\ control = Ct\ GOI\ from\ the\ Archean - Ct\ HG\ from\ the\ Archean$$

$$\Delta Ct\ test\ sample = Ct\ GOI\ from\ eCO2\ or\ PAL - Ct\ HG\ from\ HC\ or\ LC$$

Then the ΔCt values of the experimental test samples were adjusted to the ΔCt values of the control samples

$$\Delta\Delta C_t = \Delta C_{t\ test\ sample} - \Delta C_{t\ control}$$

The fold difference of the control was calculated using the following equation-

$$Fold\ change\ Control = 2\text{\^{}} - \Delta C_t$$

The fold difference of the relative gene expression of test conditions (ΔΔCt) were calculated using the following equation-

$$Fold\ change\ test = 2\text{\^{}} - \Delta\Delta C_t$$

### Supplementary Figures

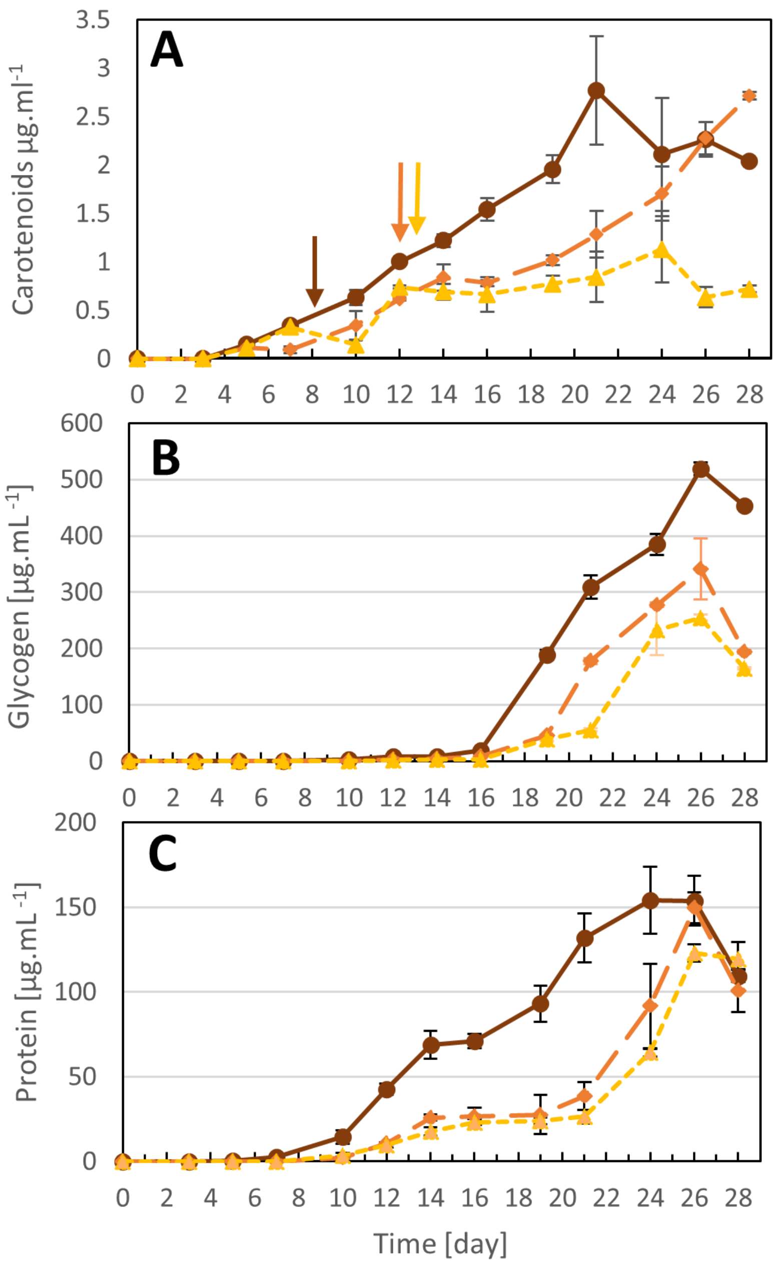


**Supplementary Figure 1. Growth assessment of *Pseudanabaena* sp. PCC7367 under three different atmospheric conditions.** Cellular carotenoid (A), glycogen (B) and protein (C) content were monitored over 28 days. Glycogen and protein content were below the level of detection during early exponential phase and are plotted as zero. Samples were taken from unstirred cultures for RNA extraction and enzyme activity determinations on day 8-9 under Archean atmospheric conditions (●), day 12-13 for the high $CO_2$ atmosphere grown cultures (◆) and day 13-14 for present-day atmospheric conditions (▲), as indicated by the arrows (A). Points present the values for three biological replicates with the bars illustrating the standard deviation. Pairwise multiple comparison (Tukey Test) indicates a significantly higher carotenoid content (to day 24) as well as glycogen and protein when compared to HC and LC grown cultures ($p<0.05$).

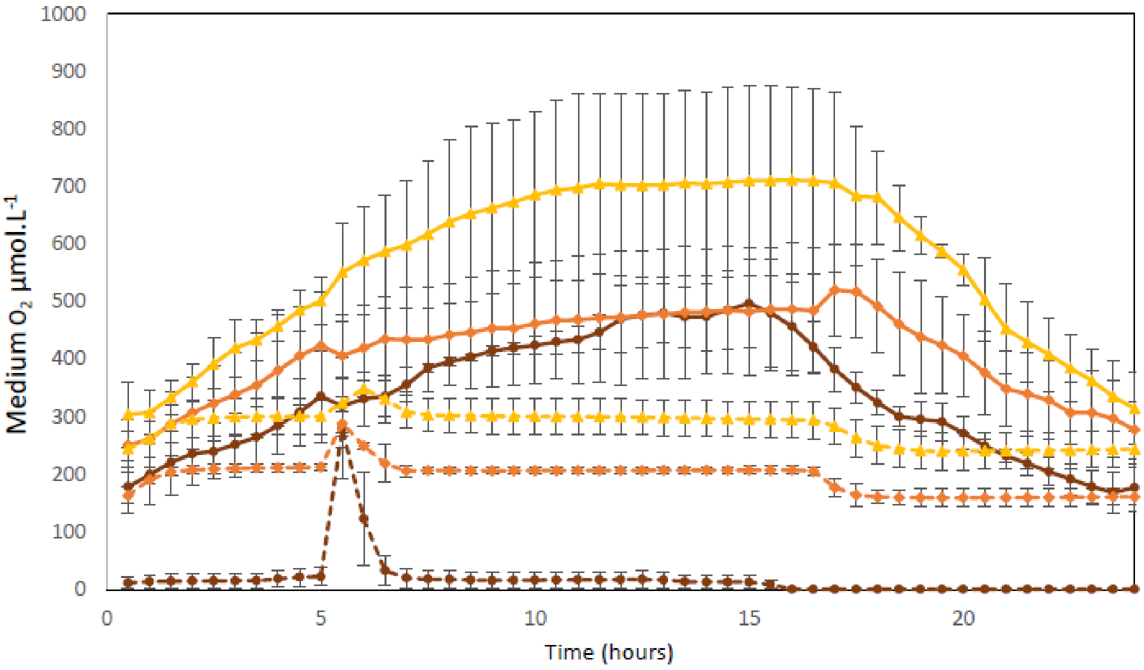


**Supplementary Figure 2. Medium $O_2$ concentration monitored for the 48 hr period of sampling for RNA extraction and enzyme activity assays**. The oxygen sensors were placed inside the unstirred cultures and monitored for 24 hours (solid lines) through day 8-9 under Archean atmospheric conditions (●), day 12-13 for the high $CO_2$ atmosphere grown cultures (◆) and day 13-14 for normal atmospheric conditions (▲). The cultures were subsequently stirred (dotted lines) and the $O_2$ levels monitored for another 24 hours through day 9-10 under Archean atmospheric conditions (●), day 13-14 for the high $CO_2$ atmosphere grown cultures (◆) and day 14-15 for normal atmospheric conditions (▲). Points represent the average of three biological replicates, averaged every 30 minutes. Bars indicate the standard deviation.

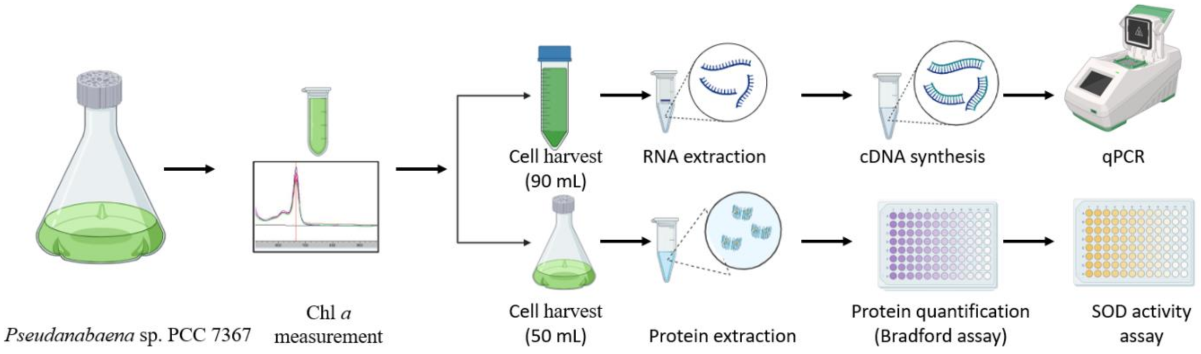


**Supplementary Figure 3: Workflow for SOD gene expression analysis and enzyme activity determination in *Pseudanabaena* sp. PCC7367.** *Pseudanabaena* sp. PCC7367 was cultivated under three different atmospheric conditions: anoxic Archean atmospheric conditions, atmosphere with a high $CO_2$ (HC) and present-day atmospheric conditions (LC), with three replicates for each atmospheric condition (n = 3). The cells were grown until the late exponential phase, determined by Chlorophyll *a* (Chl *a*) measurements. Once a similar Chl *a* concentration of approximately 2 µg/mL was reached (on day 8-9 for anoxic Archean, day 12-13 for HC, and day 13-14 for LC), cells were harvested at five distinct time points for both RNA extraction and enzyme activity determinations.

RNA extraction was performed using a 90 mL culture, and the extracted RNA was used for cDNA synthesis (mentioned in Materials and Methods). The synthesized cDNA was then utilized for gene expression analysis.

For the determination of SOD enzyme activity, protein extraction was carried out from the 50 mL culture, quantified using the Bradford assay. An equivalent amount of protein was then used for the quantification of SOD enzyme activity, as detailed in the Materials and Methods section.

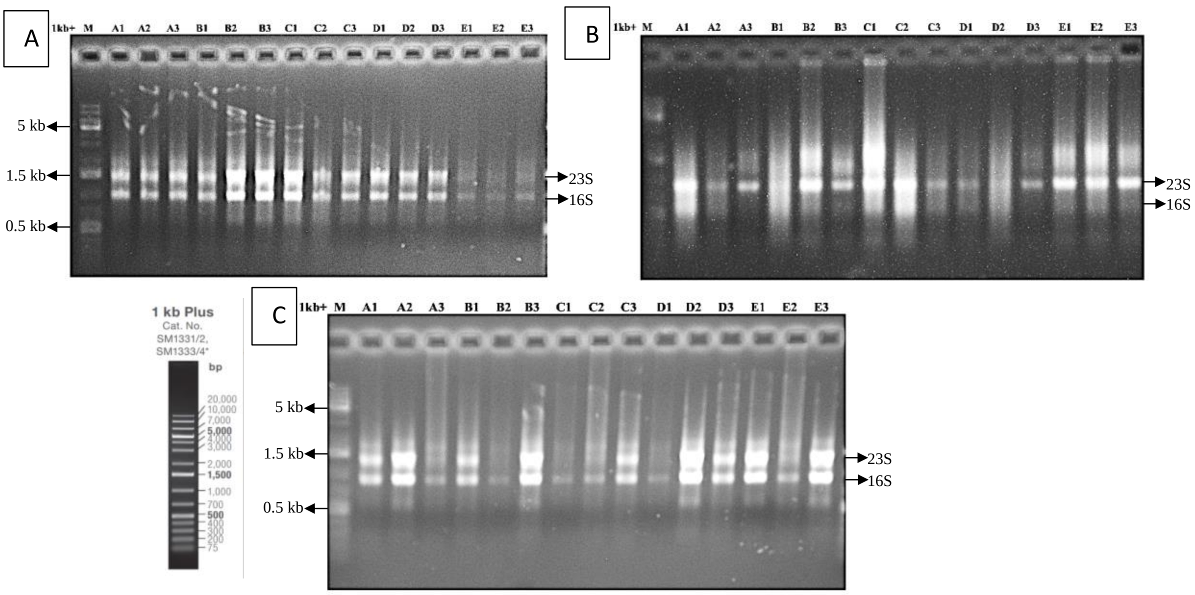


**Supplementary Figure 4: RNA isolated from *Pseudanabaena* sp. PCC 7367 growing under different atmospheric conditions.** RNA was extracted from cells grown under the Archean simulated atmospheres (A), HC atmosphere (B) and LC atmosphere (C) for 5 timepoints and visualised on an ethidium bromide stained 1 % TAE agarose gel. The three major bands representing 23 S, 16 S and 5 S RNA are visible on the gel. The smeared bands are partly degraded RNA. Samples are numbered A – E to represent the 5 different time points samples, 2 (A), 14 (B), 18 (C) and 22 (D) hours under stationary conditions and 19 (E) hours under stirred conditions, respectively (n=3).

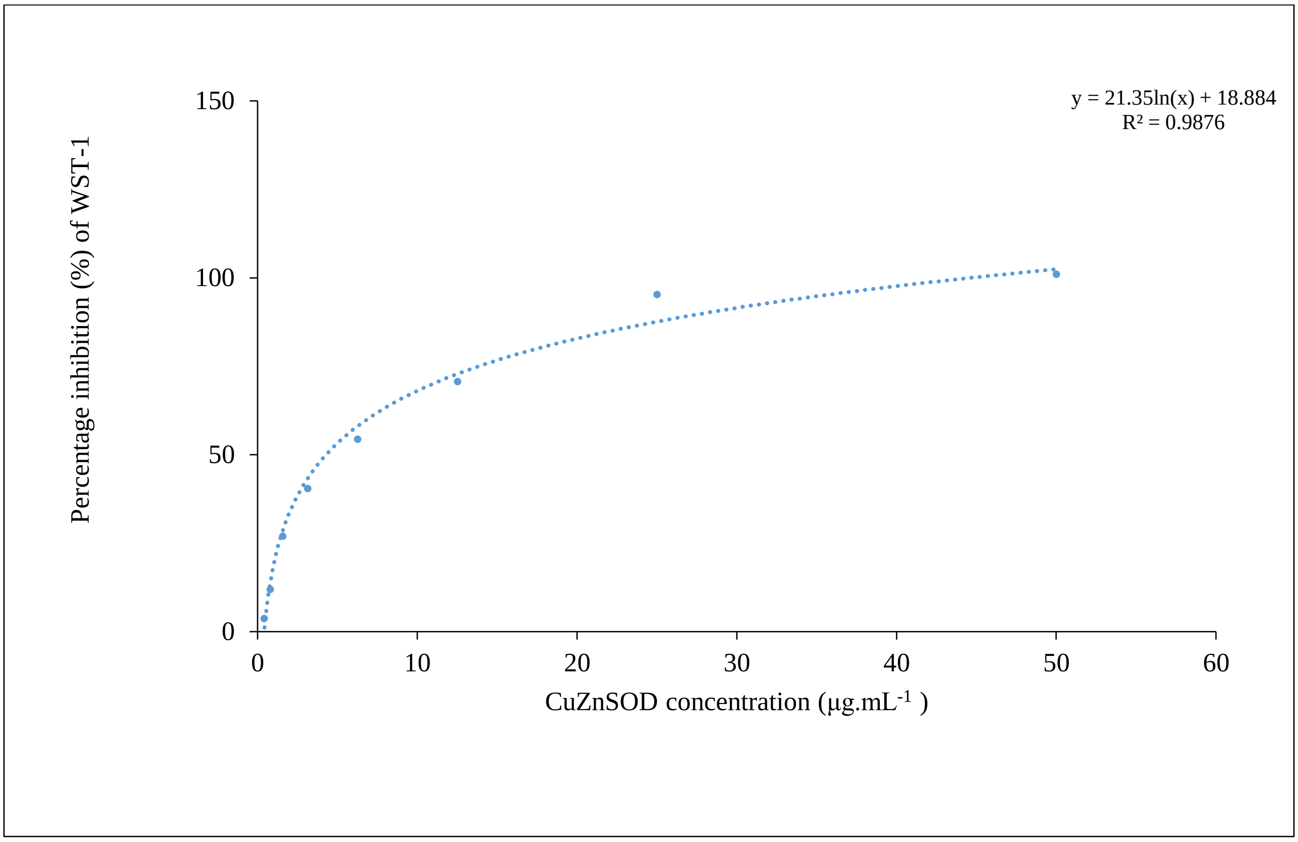


**Supplementary Figure 5: An example of a SOD standard curve using bovine CuZnSOD as standard.** Percentage inhibition (%) of WST-1 reduction by SOD is plotted against known concentrations of the CuZnSOD standard control. From the logarithmic trendline an equation was generated for CuZn SOD protein concentration (X) against percent of inhibition (Y).

**A**

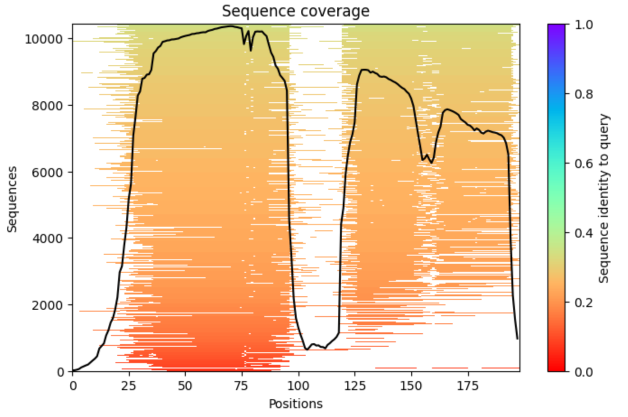


**B**

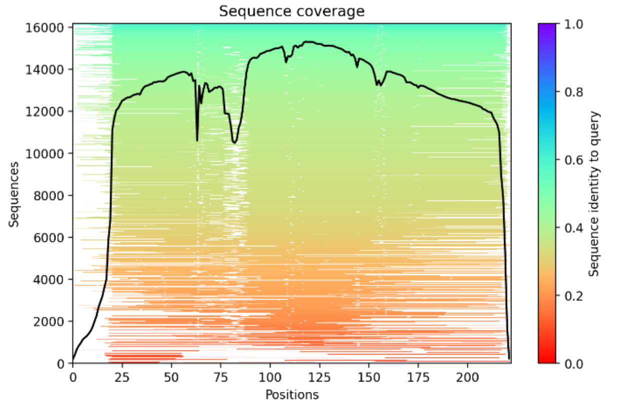


**C**

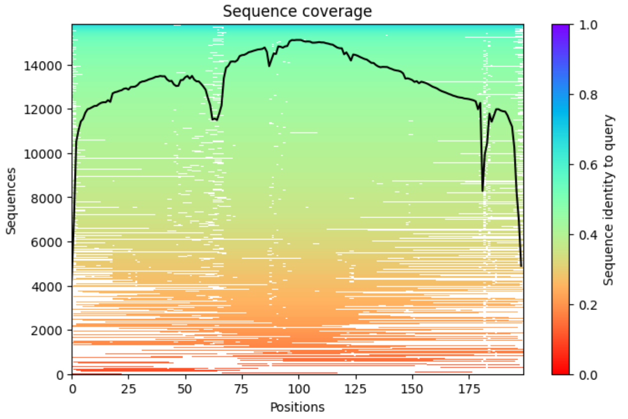


**Supplementary Figure 6**: Graphical representation of the multiple sequence alignment (msa) used to generate the Alphafold2 predicted structures in Fig. 5 for the truncated CuZnSOD (A) and MnSOD (B), as well as FeSOD (C). Shading ranges from dark blue for 100% sequence identity to bright red representing no identity to the query sequence.
