## Supplementary tables for "Early-branching cyanobacteria up-regulate superoxide dismutase activity under a simulated early Earth anoxic atmosphere"

**Supplementary Table 1: Primer sequences.** Primers specific for the superoxide dismutase genes, *sodA* (MnSOD), *sodB* (FeSOD) and *sodC* (Cu/ZnSOD), identified in *Pseudanabaena* sp. PCC7367 were designed as indicated below, as well as the reference gene primer pair targeting the *rpoC1* gene encoding the RNA polymerase beta subunit (Enzingmüller-Bleyl et al., 2022).

| Gene of interest | Region in genome CP003592.1 | Corresponding protein (NCBI Accession) | Locus | Primer direction | Sequence (5´ → 3´) | Tm [°C] | GC content (%) | Product length in base pairs (bp) |
|---|---|---|---|---|---|---|---|---|
| *sodA* | 770,159 to 770,920 | WP_015163874.1 | Pse7367_0596 | Forward | GCGCTCATGCCTGCTAAATC | 60.04 | 55 | 167 |
| | | | | Reverse | CCTGATGACCCGTTTACGCT | 60.11 | 55 | |
| *sodB* | 3,547,029 to 3,547,598 | WP_015166022.1 | Pse7367_2813 | Forward | ACTGGCGCTTTGGCTAGTAA | 57.3 | 50 | 165 |
| | | | | Reverse | GGGATTTTCAGCGTTGCCAG | 60.11 | 56.4 | |
| *sodC* | 505,866 to 506,558 | WP_015163683.1 | Pse7367_0398 | Forward | GCTGCTATGGGAGTTGTGGT | 60.04 | 55 | 198 |
| | | | | Reverse | TTGGAGGTGATCGTTGAGGC | 60.04 | 55 | |
| | | | | 809R | GCTTCGGCACGGCTCGGGTCGATA | 69,5 | 66,7 | |
| *rpoC1* | n/a | n/a | Pse7367_0455 | Forward | TGTTGGGTAAACGGGTTGAC | 57.3 | 50 | 136 |
| | | | | Reverse | CGAATCAGGCGATTAATCACAAA | 57.31 | 39.1 | |

**Supplementary Table 2: Primer binding efficiencies.** The binding efficiency of the primers designed to detect the SOD genes were determined for both genomic DNA (gDNA) and copy DNA (cDNA) reverse transcribed from RNA extracted from *Pseudanabaena* sp. PCC7367.

Genes targeted for analysis are the *sodA* (MnSOD), *sodB* (FeSOD) and *sodC* (Cu/ZnSOD) genes with the *rpoC1* gene as reference gene (Enzingmüller-Bleyl et al., 2022). The primer efficiencies were determined using genomic DNA of *Pseudanabaena* sp. PCC 7367 ranging from 10 000 000 copies to 1 copy of the targeted gene, as well as dilutions of cDNA. All genes targeted were present in a single copy on the genome of *Pseudanabaena* sp. PCC 7367.

| Gene of Interest | gDNA Dilutions | cDNA Dilutions | Primer Efficiency % with gDNA | Primer Efficiency % with cDNA |
|---|---|---|---|---|
| *rpoC1* | 10 ng | 10 ng | 107.70 | 102.68 |
| *sodA* | 10 ng | 10 ng | 95.79 | 88.7 |
| *sodB* | 10 ng | 10 ng | 89.30 | 88.7 |
| *sodC* | 10 ng | 10 ng | 98.02 | 97.88 |

**Supplementary Table 3: Secretory pathways encoded in *Pseudanabaena* sp. PCC7367.** The presence of genes encoding putative ***sec*** (Russo & Zedler, 2021; Supplementary Tables S2 & S3) and ***tat*** (Russo & Zedler, 2021) (Supplementary Table S4) pathways were obtained through a similarity search using the characterised protein sequences obtained from *E. coli* K12 MI1665 or *Synechocystis* sp. PCC6803.

| ***Gene name*** | **Bait organism** | **Bait gene** | ***Pseudanabaena* PCC 7367 uniprotkb** | **BLASTp E value** |
|---|---|---|---|---|
| *secA* | *E. coli* K12 MI1665 | P10408\|SecA | K9SFH1 | 6.81e-63 |
| *secD* | *E. coli* K12 MI1665 | P0AG90\|SecD | K9SFP1 | 1.34e-61 |
| *secF* | *E. coli* K12 MI1665 | P0AG93\|SecF | K9SFP1 | 1.81e-06 |
| *secF* | *E. coli* K12 MI1665 | P0AG93\|SecF | K9SEI8 | 2.52e-31 |
| *secY* | *E. coli* K12 MI1665 | P0AGA2\|SecY | K9SHY3 | 1.44e-107 |
| *secE* | *Synechocystis sp. PCC6803* | P38382\|SecE | K9SDF3 | 1.46e-16 |
| *secG* | *Synechocystis sp. PCC6803* | P74508\|SecG | K9SFY9 | 5.96e-21 |
| *tatA* | *E. coli* K12 MI1665 | P69428\|TatA | K9SKB5 | 2.24e-05 |
| *tatA* | *E. coli* K12 MI1665 | P69428\|TatA | K9SI06 | 5.16e-05 |
| *tatC* | *E. coli* K12 MI1665 | P69423\|TatC | K9SCK0 | 2.55e-27 |

**Supplementary Table 4: Metal transporters in *Pseudanabaena* sp. PCC7367.** The presence of metal transporters encoded by *Synechocystis* PCC6803 for the metal cofactors required for the SOD isoforms investigated in this study, is indicated in the table below (modified from Sharon *et al.*, 2014). The characterized protein sequences from *Synechocystis* sp. PCC6803 (NC_000911.1) were used to search for similarity homologues in the genome of *Pseudanabaena* sp. PCC7367 (NC_019701.1) using tblastn (Altschul, 1991).

| **Metal** | ***Gene name*** | **Description** | ***Synechocystis* 6803 NC_000911.1 Gene locus tag** | ***Synechocystis* 6803 Protein ID** | ***Pseudanabaena* PCC 7367 NC_019701.1 Gene locus tag** | **Description** | **BLASTp E value** |
|---|---|---|---|---|---|---|---|
| **Mn** | *mntC* | PsaA, metal ABC transporter substrate binding domain | SGL_RS08820 | WP_010872546.1 | WP_015163932.1 | zinc ABC transporter substrate-binding protein | 4e-44 |
| | *mntA* | Inorganic $Mn^{2+}/Zn^{2+\ +}$ ion transporter ATPase component | SGL_RS08815 | WP_010872545.1 | WP_015163931.1 | metal ABC transporter ATP-binding protein | 8e-59 |
| | *mntB* | ABC 3 transporter permease | SGL_RS0881 | WP_010872544.1 | WP_015166116.1 | ABC-type $Mn^{2+}/Zn^{2+}$ transport system, permease | 1e-59 |
| **Zn** | *znuB* | ABC-type $Mn^{2+}/Zn^{2+}$ transport system, ATP-binding protein | SGL_RS04625 | WP_010871740.1 | WP_015163931.1 | ABC-type $Mn^{2+}/Zn^{2+}$ transport system, ATPase | 1e-43 |
| | *znuC* | hypothetical | SGL_RS04630 | WP_010871742.1 | / | | No hit |
| | *znuB (ZntC?)* | zinc ABC transporter substrate-binding protein | SGL_RS04620 | WP_010871739.1 | WP_015166551.1 | zinc ABC transporter substrate-binding protein | 3e-28 |
| **Cu** | *ctaA* | putative Cu binding site & ATP binding site | SGL_RS12475 | WP_010873238.1 | WP_051038173.1 | copper-translocating P-type ATPase | 0.0 |
| | *pacS* | copper-translocating P-type ATPase | SGL_RS05435 | WP_010871897.1 | WP_015164254.1 | copper-translocating P-type ATPase | 0.0 |

**Supplementary Table 5: Potential genes involved in the reduction of $H_2O_2$, the product of SOD dismutation of $O_2^{-\bullet}$ .**

The presence of genes potentially encoding peroxidases or peroxiredoxins were obtained from the KEGG database (Kanehisa et al., 2022) for the genome of *Pseudanabaena* sp. PCC7367 (NC_019701.1). The presence of a glutathione synthase and glutathione reductase were also confirmed.

| **Description** | ***Pseudanabaena* sp. PCC7367 KEGG entry** | **Description** | **BLAST against the protein data bank and manual sequence inspection** |
|---|---|---|---|
| Peroxiredoxin | Pse7367_1682 | thioredoxin-dependent peroxiredoxin | BCP-type enzyme (alternative reductants) |
| Peroxiredoxin | Pse7367_2013 | thioredoxin-dependent peroxiredoxin | Prx1-type enzyme (thioredoxin as a reductant) |
| Peroxiredoxin | Pse7367_2293 | thioredoxin-dependent peroxiredoxin | BCP-type enzyme (alternative reductants) |
| Redoxin domain protein | Pse7367_2374 | thioredoxin-dependent peroxiredoxin | Prx5-type enzyme (alternative reductants) |
| 1-Cys peroxiredoxin | Pse7367_3020 | thioredoxin-dependent peroxiredoxin | Prx6-type enzyme (reductant unknown) |
| Oxidoreductases | Pse7367_2328 | peroxidase | Heme-containing peroxidase |
| Glutathione peroxidase | Pse7367_3393 | glutathione peroxidase | Trx-dependent GPx (thioredoxin as a reductant) |
| Glutathione synthase | Pse7367_3280 | | |
| Glutathione reductase | Pse7367_0547 | NADPH-glutathione reductase | |

**Supplementary Table 6:** Results of One-way repeated measure ANOVA and/or ANOVA on ranks with environmental treatment condition as factor and Post-Hoc all pairwise multiple comparison procedures using Turkey Test and/or Holm-Sidak method for biomass parameters (Fig. 2 Suppl Fig 1). Bold numbers indicate significant differences with minimum p≤0.05. Number of replicates = 3.

| **One-Way repeated measure biomass parameter** | | | | | | |
|---|---|---|---|---|---|---|
| | **One-way repeated measure ANOVA** | | | **ANOVA on ranks** | | |
| | **F** | **p** | **DF** | **Chi-square** | **p** | **Df** |
| **Chlorophyll a** | | | | **22,00** | **≤0.001** | **2** |
| **Carotenoid** | **8,443** | **0,002** | **2** | | | |
| **Protein** | | | | **15.048** | **≤0.001** | **2** |
| **Glycogen** | **7,01** | **0,004** | **2** | | | |
| **Growth** | **214,102** | **≤0.001** | **2** | | | |

| **Post-Hoc pairwise multiple comparison** | | | | | | |
|---|---|---|---|---|---|---|
| | **LC vs. Archean** | | **LC vs. HC** | | **HC vs. Archean** | |
| | *q/t* | *p* | *q/t* | *p* | *q/t* | *p* |
| **Chlorophyll a** | **6.351** | **<0.001** | 3.175 | 0.064 | 3.175 | 0.064 |
| **Carotenoide** | **5.804** | **0.001** | 3.15 | 0.087 | 2.655 | 0.167 |
| **Protein** | **4.715** | **0.002** | 1.109 | 0.713 | **3.606** | **0.029** |
| **Glycogen** | **5.103** | **0.004** | 1.328 | 0.622 | **3.776** | **0.035** |
| **Growth** | **18.930** | **<0.001** | 2.227 | 0.068 | **16.703** | **<0.001** |

**Supplementary Table 7:** Results of One-way repeated measure ANOVA with environmental treatment condition as factor for growth media oxygen concentration (Suppl. Fig.2). In case of significant differences, Post-Hoc analysis using Holm-Sidak method was applied to identify differing groups: Bold numbers indicate significant differences with minimum p≤0.05. Number of replicates = 3.

| **One-Way repeated measure for growth media oxygen concentration** | | | |
|---|---|---|---|
| **Time after onset of monitoring** | **One-way repeated measure ANOVA** | | |
| | **F** | ***p*** | **DF** |
| **2h** | 3.399 | 0.137 | **2** |
| **14h** | 2.826 | 0.172 | **2** |
| **18h** | **17.941** | **0.01** | **2** |
| **22h** | **7.411** | **0.045** | **2** |
| **19h stirred** | **125.449** | **<0.001** | **2** |

| **Post-Hoc pairwise multiple comparison** | | | | | | |
|---|---|---|---|---|---|---|
| **Time after onset of monitoring** | **LC vs. Archean** | | **LC vs. HC** | | **HC vs. Archean** | |
| | *t* | *p* | *t* | *p* | *t* | *p* |
| **2h** | n.s. | | | | | |
| **14h** | n.s. | | | | | |
| **18h** | **5.988** | **0.012** | **2.872** | **0.045** | 3.117 | 0.07 |
| **22h** | **5.401** | **0.04** | 2.102 | 0.388 | 3.299 | 0.161 |
| **19h stirred** | **15.575** | **<0.001** | **5.289** | **0.006** | **10.286** | **0.001** |

**Supplementary Table 8 :** Results of repeated measure two way ANOVA using environmental treatment condition (Arch, HC, LC) as well as sampling time (Fig. 3, A-D) as factor for SOD gene expression, SOD protein activity and dissolved $O_2$ in the growth media. In case of significant differences, pairwise multiple comparison procedure using Duncan's Method for treatment (A) and time (B) as factor was applied to identify the respective groups of parameters that differ significantly. Bold numbers indicate significant differences with minimum p≤0.05. Number of replicates = 3.

| | **Treatment** | | **Time** | | **Treatment x Time** | |
|---|---|---|---|---|---|---|
| | *F-value* | *p* | *F-value* | *p* | *F-value* | *p* |
| **qPCR *sodA* (MnSOD)** | 5,156 | 0,078 | **9,183** | **0,004** | **24,669** | **<0.001** |
| **qPCR *sodB* (FeSOD)** | **105,979** | **<0.001** | **61,534** | **<0.001** | **27,317** | **<0.001** |
| **qPCR *sodC* (CuZnSOD)** | **125,71** | **<0.001** | **21,583** | **<0.001** | **6,632** | **<0.001** |
| **qPRC *sodABC* Sum** | **102,919** | **<0.001** | **22,448** | **<0.001** | **4,189** | **0,007** |
| **SOD enzyme activity** | **85,075** | **<0.001** | **540,712** | **<0.001** | **158,534** | **<0.001** |
| **Dissolved $O_2$** | **11,069** | **0,023** | **112,171** | **<0.001** | **3,167** | **0,024** |
| **Post-Hoc pairwise multiple comparison** | | | | | | |
| **A) Factor Treatment** | | | | | | |
| | **LC vs. Archean** | | **LC vs. HC** | | **HC vs. Archean** | |
| | *q* | *p* | *q* | *p* | *q* | *p* |
| **qPCR *sodA* (MnSOD)** | **4,534** | **0,035** | 2,486 | 0,154 | 2,048 | 0,221 |
| **qPCR *sodB* (FeSOD)** | **16,783** | **<0.001** | 1,938 | 0,243 | **18,721** | **<0.001** |
| **qPCR *sodC* (CuZnSOD)** | **22,423** | **<0.001** | **11,361** | **0,002** | **11,063** | **0,002** |
| **qPRC *sodABC* Sum** | **18,704** | **<0.001** | 2,543 | 0,147 | **16,162** | **<0.001** |
| **SOD enzyme activity** | **17,118** | **<0.001** | 2,603 | 0,14 | **14,514** | **<0.001** |
| **Dissolved $O_2$** | **6,652** | **0,01** | 3,484 | 0,07 | 3,167 | 0,089 |
| **B) Factor Time** | | | | | | |
| | **qPCR *sodA* (MnSOD)** | | **qPCR *sodB* (FeSOD)** | | **qPCR *sodC* (CuZnSOD)** | |
| | *q* | *p* | *q* | *p* | *q* | *p* |
| 14.000 vs. 18.000 | 0.0145 | 0.992 | 12.15 | **<0.001** | 0.187 | 0.898 |
| 14.000 vs. 19.000 | 4.041 | **0.025** | 19.701 | **<0.001** | 11.234 | **<0.001** |
| 14.000 vs. 2.000 | 0.0272 | 0.985 | 17.263 | **<0.001** | 4.438 | **0.019** |
| 14.000 vs. 22.000 | 4.521 | **0.015** | 16.668 | **<0.001** | 1.943 | 0.224 |

| | | | | | | |
|---|---|---|---|---|---|---|
| 18.000 vs. 19.000 | 4.056 | **0.028** | 7.552 | **0.001** | 11.047 | **<0.001** |
| 18.000 vs. 2.000 | 0.0416 | 0.979 | 5.113 | **0.008** | 4.251 | **0.02** |
| 18.000 vs. 22.000 | 4.507 | **0.013** | 4.519 | **0.013** | 1.756 | 0.25 |
| 2.000 vs. 19.000 | 4.014 | **0.022** | 2.439 | 0.123 | 6.796 | **0.002** |
| 22.000 vs. 19.000 | 8.563 | **<0.001** | 3.033 | 0.074 | 9.291 | **<0.001** |
| 22.000 vs. 2.000 | 4.549 | **0.017** | 0.594 | 0.685 | 2.495 | 0.116 |
| | **qPRC *sodABC* Sum** | | **SOD enzyme activity** | | **Dissolved $O_2$** | |
| | *q* | *p* | *q* | *p* | *q* | *p* |
| 14.000 vs. 18.000 | 0.0145 | 0.992 | 12.15 | **<0.001** | 0.187 | 0.898 |
| 14.000 vs. 19.000 | 4.041 | **0.025** | 19.701 | **<0.001** | 11.234 | **<0.001** |
| 14.000 vs. 2.000 | 0.0272 | 0.985 | 17.263 | **<0.001** | 4.438 | **0.019** |
| 14.000 vs. 22.000 | 4.521 | **0.015** | 16.668 | **<0.001** | 1.943 | 0.224 |
| 18.000 vs. 19.000 | 4.056 | **0.028** | 7.552 | **0.001** | 11.047 | **<0.001** |
| 18.000 vs. 2.000 | 0.0416 | 0.979 | 5.113 | **0.008** | 4.251 | **0.02** |
| 18.000 vs. 22.000 | 4.507 | **0.013** | 4.519 | **0.013** | 1.756 | 0.25 |
| 2.000 vs. 19.000 | 4.014 | **0.022** | 2.439 | 0.123 | 6.796 | **0.002** |
| 22.000 vs. 19.000 | 8.563 | **<0.001** | 3.033 | 0.074 | 9.291 | **<0.001** |
| 22.000 vs. 2.000 | 4.549 | **0.017** | 0.594 | 0.685 | 2.495 | 0.116 |

**Supplementary Table 9:** Pearson's correlation of parameters monitored during 24 h sampling (Fig. 3). Bold numbers indicate significant differences with minimum p≤0.05. Abrev. r= correlation coefficient, n= number of samples. Number of replicates = 3.

| | | **qPCR *sodA*** | **qPCR *sodB*** | **qPCR *sodC*** | **qPCR *sodABC* sum** | **SOD enzyme activity** | **medium $O_2$** |
|---|---|---|---|---|---|---|---|
| **Time** | r | 0.0796 | -0.0238 | -0.00657 | 0.0209 | -0.0844 | -0.0417 |
| | *p* | 0.603 | 0.877 | 0.966 | 0.892 | 0.582 | 0.785 |
| | n | 45 | 45 | 45 | 45 | 45 | 45 |
| **qPCR *sodA*** | r | | -0.149 | **0.521** | **0.47** | **0.349** | 0.234 |
| | *p* | | 0.327 | **0.000242** | **0.00114** | **0.0186** | 0.122 |
| | n | | 45 | 45 | 45 | 45 | 45 |
| **qPCR *sodB*** | r | | | **0.382** | **0.797** | 0.071 | **0.651** |
| | *p* | | | **0.00958** | **5.71E-11** | 0.643 | **0.00000127** |
| | n | | | 45 | 45 | 45 | 45 |
| **qPCR *sodC*** | r | | | | **0.725** | 0.0142 | **0.675** |
| | *p* | | | | **1.79E-08** | 0.926 | **0.000000376** |
| | n | | | | 45 | 45 | 45 |
| **qPCR *sodABC* sum** | r | | | | | 0.242 | **0.751** |
| | *p* | | | | | 0.11 | **2.81E-09** |
| | n | | | | | 45 | 45 |
| **SOD enzyme activity** | r | | | | | | 0.202 |
| | *p* | | | | | | 0.183 |
| | n | | | | | | 45 |

References:

Altschul, S. F. (1991). Amino acid substitution matrices from an information theoretic perspective. *Journal of molecular biology*, *219*(3), 555-565.

Enzingmüller-Bleyl, T. C., Boden, J. S., Herrmann, A. J., Ebel, K. W., Sánchez-Baracaldo, P., Frankenberg-Dinkel, N., & Gehringer, M. M. (2022). On the trail of iron uptake in ancestral Cyanobacteria on early Earth. *Geobiology*.

Kanehisa, M., Furumichi, M., Sato, Y., Kawashima, M., & Ishiguro-Watanabe, M. (2022). KEGG for taxonomy-based analysis of pathways and genomes. *Nucleic acids research*, *51*(D1), D587-D592. https://doi.org/10.1093/nar/gkac963

Russo, D. A., & Zedler, J. A. (2021). Genomic insights into cyanobacterial protein translocation systems. *Biological chemistry, 402*(1), 39-54.
